# A handheld microfluidic manifold for massively multiplexed nucleic acid detection

**DOI:** 10.1101/2025.08.11.669785

**Authors:** Sijbren Kramer, Ryungeun Song, Yujia Huang, Soonwoo Hong, Ibrahim Motlani, Howard A. Stone, Cameron Myhrvold

## Abstract

Multiplexed methods for nucleic acid detection are immensely challenging to deploy outside of laboratory settings. Conversely, field-deployable methods are limited to low levels of multiplexing. Here, we introduce Scalable On-site Nucleic Acid Testing Architecture (SONATA), enabling >100-chamber reaction partitioning for multiplexed nucleic acid amplification and detection with a portable microfluidic manifold. The manifold directs a diluted sample into individual reaction chambers, each of which contains lyophilized reagents and a small stir bar or bead for mixing. Samples can be loaded using a syringe by hand, greatly simplifying the testing process. We demonstrate the integration of the platform with Streamlined Highlighting of Infections to Navigate Epidemics (SHINE), a sensitive and deployable CRISPR-based detection technology. We show that deployed with SONATA, SHINE retains its sensitivity, enabling highly multiplexed pathogen detection in ≤ 1 hour. In addition, we demonstrate the detection of single-nucleotide variants, including mutations associated with drug susceptibility.

## Introduction

Multiplexed nucleic acid detection is critical for the surveillance of pathogens (*1*–*3*), cancer mutations (*4, 5*), and genetic diseases (*6, 7*). However, existing technologies either require expensive laboratory equipment and specially-trained personnel, such as next-generation sequencing (NGS) and polymerase chain reaction (PCR), or are limited to low levels of multiplexing, like isothermal amplification (*8, 9*). This makes the routine monitoring of a wide array of pathogens and mutations infeasible, particularly in resource-limited settings. An ideal surveillance technology would combine the high multiplexing capacity of NGS with the simplicity, speed, and low cost of isothermal nucleic acid amplification tests to ultimately allow for distributed multiplexed testing that can be performed by anyone, anywhere.

Certain technologies attempt to bridge this gap. Coupling PCR with portable sequencing platforms such as the Oxford Nanopore MinION enables extremely high multiplexing through amplicon sequencing and barcoding (*10*), but demands substantial library preparation steps and bioinformatics infrastructure, complicating deployment outside well-equipped labs (*11*). Integrated cartridge-based PCR systems such as BioFire FilmArray combine automated sample preparation and nested multiplex PCR to achieve high multiplexing (>30-plex) (*12*). Nonetheless, the reliance on sophisticated and expensive laboratory equipment (>$30,000) limits its use in resource-limited settings (*13*). In contrast, isothermal amplification methods including loop-mediated isothermal amplification (LAMP) and recombinase polymerase amplification (RPA), which eliminate thermal cycling, can be deployed with low-cost platforms such as paper microfluidics. However, to date, these approaches have enabled multiplexed detection of only a single-digit number of targets (*14*).

In recent years, deployable diagnostic assays leveraging CRISPR have emerged that combine the sensitivity of isothermal amplification with the specificity of CRISPR effector proteins, Cas12 and Cas13 (*8, 15*–*22*). Simultaneous detection of 2-4 targets has been achieved in single-pot reactions using orthogonally-cleaving Cas proteins, but this approach is constrained by the small number of available orthogonal effectors, so it is challenging to scale beyond 4-plex (*18, 23, 24*). To overcome this limitation, microfluidic platforms have been developed to enable higher degrees of spatial multiplexing (*25*).

CARMEN (Combinatorial Arrayed Reactions for Multiplexed Evaluation of Nucleic acids) and its microfluidic version (mCARMEN) achieve highly multiplexed detection of 169-plex and 96-plex, respectively, but require extensive laboratory infrastructure (*1, 26*). MiCaR (<u>mi</u>crofluidic device with CRISPR-<u>C</u>as12a <u>a</u>nd multiplex <u>r</u>ecombinase polymerase amplification) employs isothermal amplification with up to 30-chamber reaction partitioning; however, pooled off-chip amplification limits detection to nine targets, and the need for complex liquid handling and non-lyophilized reagents restricts field deployability (*27*). Hence, there remains a need for a deployable highly-multiplexed nucleic acid detection technology.

Here, we introduce Scalable On-site Nucleic Acid Testing Architecture (SONATA), a microfluidic platform that deploys nucleic acid amplification in a highly multiplexed manner (up to 112-chamber reaction partitioning). SONATA utilizes spatial multiplexing, in which a custom microfluidic manifold distributes a sample fluid and resuspension buffer mixture to separate reaction chambers pre-loaded with lyophilized reagents. The design incorporates droplet lyophilization, magnetic on-chip mixing, reaction chamber oil/air sealing, and a fluorescent visual readout. We demonstrate the sensitivity, specificity, and multiplexing capability of SONATA by deploying the one-pot isothermal CRISPR-based nucleic acid detection technology, Streamlined Highlighting of Infections to Navigate Epidemics (SHINE), on Mpox, *Mycobacterium tuberculosis* (Mtb), and epidermal growth factor receptor (EGFR) mutations implicated in non-small cell lung cancer (NSCLC) drug response.

## Results

As shown in **Fig. 1A**, a syringe is used to manually inject a mixture of sample fluid and resuspension buffer into the chip. This injection process takes less than a minute. The chip geometry automatically distributes flow evenly to reaction chambers with pre-loaded lyophilized reagents. Any excess fluid is contained in an on-chip reservoir, preventing user contact with the sample. Next, an oil or air sealing step is performed with a squeeze bottle or syringe, fully sealing the reaction chambers, which is followed by a simple magnetic mixing process for one minute. The chip is then incubated for 30-90 minutes and the fluorescent readout is imaged with a fluorescent imaging device.

**Figure 1.**
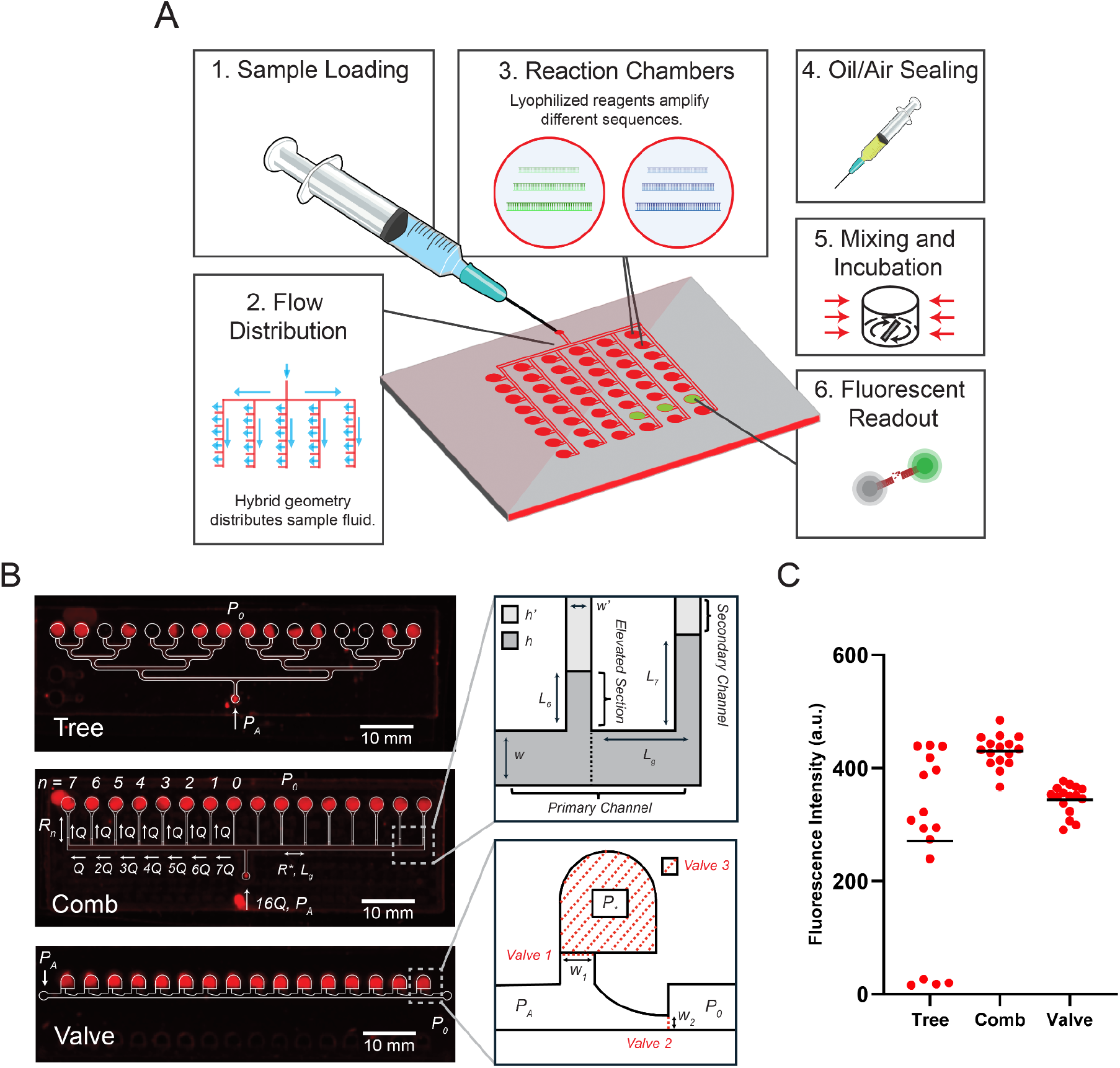
A microfluidic manifold for multiplexed nucleic acid amplification and detection. **(A)** Schematic of the SONATA workflow developed here, showing **(1)** injection of the sample and resuspension buffer mixture into the chip by a handheld syringe, **(2)** uniform flow distribution to the reaction chambers enabled by the hybrid geometry, **(3)** reaction chambers containing lyophilized reagents designed to amplify different sequences, **(4)** an oil or air sealing step to isolate independent reaction environments, **(5)** on-chip mixing enabled by movement of the chip relative to an external magnet followed by incubation, and **(6)** a fluorescent reporter or dye readout. **(B)** Experimental fluorescent image showing the distribution of ROX dye in the reaction chambers after injection into the 16-chamber tree, comb, and valve designs with the flow distribution geometries overlaid in white. *P*_*A*_ is the applied pressure to drive flow and *P*_*0*_ is the atmospheric pressure. The **Tree geometry** relies on an equal path length for even distribution of the flow along the different paths. The **Comb geometry** relies on adjusted hydraulic resistances to achieve a uniform flow distribution. *Q* is the flow rate. *L*_*n*_ is the length of the elevated section for each channel (n ∈ {0, 1, 2, 3, 4, 5, 6, 7}) used to adjust the resistance Rn of each secondary channel. R* is the resistance of each segment of the primary channel of length *L*_*g*_. In the zoomed inset, w is the width of the primary channel, w′ is the width of the secondary channel and the elevated section, h is the height of the primary channel and the elevated section, and h′ is the height of the secondary channel. The **Valve Geometry** relies on the sequential bursting of valves for flow distribution. The diagram shows Valve 1, Valve 2, and their corresponding widths W_1_ and W_2_, as well as Valve 3, which is the hydrophobic membrane at the surface of the chamber. Valve 1 bursts before Valve 2 causing the chamber to be filled; when the fluid reaches Valve 3, it encounters extremely high flow resistance, causing Valve 2 to burst and the fluid to advance to the next chamber. *P*_*\**_ represents the elevated pressure in the chamber due to the membrane impeding the escape of air. **(C)** Plot of the chamber fluorescence intensities in **B**. The mean of n=16 technical replicates is shown by the black line. The mean fluorescence is higher for the comb than the valve geometry due to a difference in the chamber aspect ratio and shape.

### Design of the flow distribution geometries

To enable the spatial multiplexing of amplification reactions, a geometry that reliably distributes flow evenly across reaction chambers is needed. For this device, we use 10.5 μl reaction volumes. This large reaction volume is advantageous for nucleic acid amplification as it facilitates detection of low copy targets, allows for an easily visible readout, and facilitates a manageable distribution of lyophilized reagents into reaction chambers. However, the large reaction volume makes reliable and efficient reaction partitioning more difficult. To keep the device spatially compact while allowing for high levels of multiplexing, we incorporate 3D reaction chambers that extend vertically relative to their feed channels, which allows for larger reaction chambers while distribution channels are low profile to minimize the dead volume. An additional constraint expected for portability of the device is that these flow distribution geometries need to accommodate the variable applied input pressure generated by manual syringe injection.

We prototyped three different flow distribution geometries, a tree, comb, and valve geometry **(Fig. 1B)**. The tree geometry relies on successive bifurcations to distribute flow. There is a relatively high hydrodynamic resistance associated with small channel dimensions, which contributes to a large pressure drop across the geometry.

Consequently, the liquid-air interfaces are temporarily pinned at the bifurcations. Given irregularities in channel dimensions due to the limited resolution of CNC milling, this design leads to inconsistent chamber filling (**Fig. 1B**,**C**).

The comb geometry relies on the principle that we can individually adjust the hydrodynamic resistance of each of the secondary channels by changing the length of an elevated section to ensure a uniform flow rate into each chamber **(Fig. 1B)**. The calculations for the length of this elevated section are explained in the chip design methods section. We find that the comb geometry accommodates manual injection, leading to uniform flow distribution in a few seconds **(Fig. 1B,C)**. However, it contributes to a considerable increase in dead volume as the number of chambers increases.

Additionally, if the volume injected is not carefully controlled, this comb geometry has no mechanism for containing excess sample fluid, which can lead to fluid leaking through the air outlets in the optical film used to seal the top of the chip, as shown in the leftmost chamber in **Fig. 1B**.

The valve geometry utilizes the successive bursting of surface-tension-regulated valves to enable flow distribution **(Fig. 1B)**. Valve-based designs have been shown to be effective for chamber filling at large scales (*28, 29*). As the liquid is injected, the interface is pinned at valve 2 and forced upward to fill the reaction chamber. When the liquid reaches the top of the reaction chamber, an air-permeable hydrophobic membrane (valve 3), which presents extremely high flow resistance, prevents further advancement. As the pressure builds, it exceeds the burst pressure at valve 2, causing the interface to de-pin and allowing the liquid to advance and fill the subsequent chamber. The order of bursting of the valves is controlled by the width of the valves as defined by the burst pressure equation, which is explained in the methods section. This geometry also allows for an air sealing or oil sealing step to fully displace the fluid from the main channel and seal off the reaction chambers. When the main channel is treated with a hydrophobic coating (Aquapel), the valve geometry leads to uniform flow distribution via its effect on the contact angles in the burst pressure equation **(Fig. 1B)**.

### Mixing

We wanted to ensure that SONATA could incorporate mixing, as RPA in particular has been shown to be sensitive to mixing (*30, 31*). Due to the small length scales, the Reynolds number of microfluidic flows is often very low. Thus, turbulence does not occur, making mixing a significant challenge in microfluidic systems. Hence, often slow, diffusion-based mixing is relied upon (*32*).

We sought to incorporate a mechanism for mixing that did not rely on electrical input or external equipment and could be induced via a simple user action. We hypothesized that the rotation of a stir bar in each reaction chamber could be induced by the manual movement of the chip in a circular motion relative to an external magnetic strip. To test this idea, we use ROX dye in a viscous 100% glycerol solution to evaluate mixing.

Qualitatively, we find that after only 30 seconds of hand-induced mixing the distribution of ROX dye appears to be uniform, and shows a significant improvement relative to reaction chambers without a mixing bar (**Fig. 2A)**. A confocal quantification of a reaction chamber shows a much more uniform distribution of ROX fluorescence after 30 seconds of mixing with the bar, whereas the 10 minute diffusion-only condition without the mixing bar shows a similar distribution to the initial spatial distribution of ROX **(Fig. 2B)**. In later experiments, we transition from using a stir bar to a stir bead for more uniformity and ease of handling.

**Figure 2.**
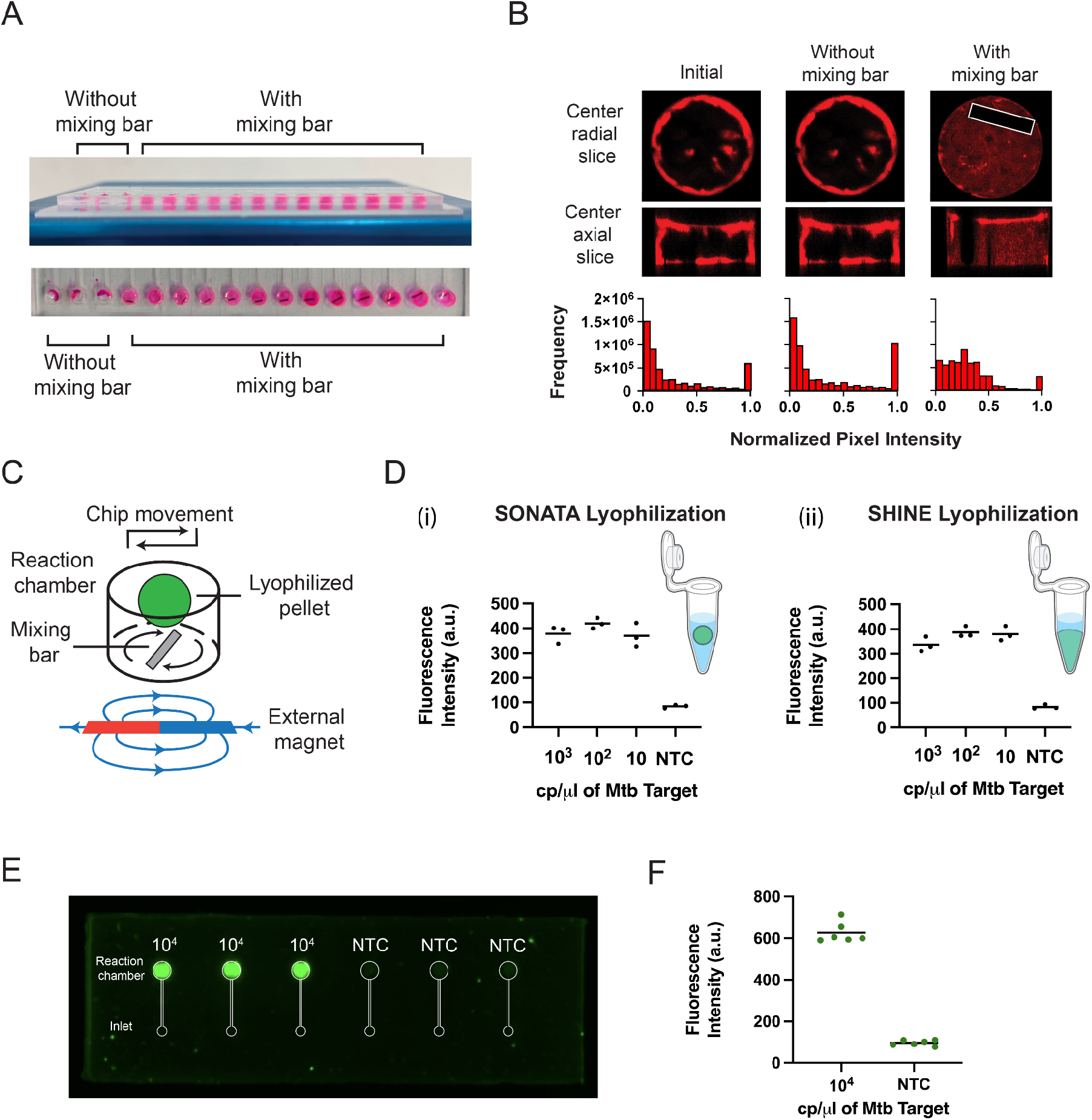
Droplet lyophilization and magnetic mixing enable deployable on-chip nucleic acid detection using SHINE. **(A**) Mixing analysis was performed with ROX dye in a viscous 100% glycerol solution to evaluate mixing. Images of the front and top of the comb chip show that after 30 seconds of mixing, the three leftmost chambers that do not contain mixing bars show an uneven distribution of ROX dye, whereas the rest of the chambers, which do contain mixing bars, appear well-mixed. **(B)** Confocal microscopy was used to generate a series of radial slices along the height dimension of the chamber. Center radial and axial slices of the reaction chamber show the distribution of ROX dye for the start condition (Initial), after a 10 minute diffusion period (Without mixing bar), and after 30 seconds of mixing with the bar (With mixing bar). The stir bar is visible in the ‘With mixing bar’ condition. Histograms show normalized pixel intensities across the entire series of confocal images. **(C)** The schematic of the reaction chamber illustrates how moving the chip by hand in a circular motion relative to a stationary external magnet induces a rotation of the mixing bar in each reaction chamber, enabling mixing of the reagents, which are pre-loaded in the form of a lyophilized pellet. **(D)** Quantification of the fluorescence of n=3 technical replicates of the SONATA and SHINE lyophilization procedures for different concentrations of Mtb targets in strip tubes showing similar limits of detection at 60 minutes. **(i)** Detection for the SONATA droplet lyophilization procedure described in the methods section. The schematic shows the more compact lyophilized pellet in the resuspension buffer. **(ii)** Detection for the SHINE lyophilization procedure described in the methods section. **(E)** FAM fluorescent image of n=3 technical replicates of the on-chip detection of 10^4^ cp/μl of Mtb synthetic DNA target and a no target control (NTC) at 60 minutes, using the SONATA lyophilization procedure and mixing approach. **(F)** Quantification of FAM fluorescent images of two chips with the black line showing the mean of n = 6 technical replicates.

### Lyophilization

We decided to integrate the Cas13-RPA SHINE technology with SONATA for further optimization and validation due SHINE’s previously demonstrated sensitivity and specificity (*22, 24*). In this one-pot reaction, DNA targets are amplified via RPA and then transcribed to synthesize RNA to enable Cas13 targeting. If Cas13’s reconfigurable crRNA binds to its specific RNA target sequence, this triggers the collateral cleavage of surrounding RNA sequences. In particular, the cleavage of a fluorescein amidite (FAM) fluorophore attached to a quencher via a polyuracil RNA sequence yields a detectable fluorescent signal, indicating the presence of the sequence of interest **(Fig. S1B**).

Often, microfluidic chips developed for nucleic acid detection have relied on liquid reagents that must be pipetted into the reaction chambers (*25*). Without on-chip storage of reagents, the liquid handling steps required to distribute reagents to reaction chambers are prohibitive to the point-of-care use of any such device. On-chip storage is central to achieving deployable multiplexing. Lyophilization (freeze drying) is the process by which a solution is frozen and then, while still frozen, exposed to a vacuum such that the water and/or other solvents are extracted via sublimation (*33*). It has been demonstrated previously that lyophilization is an effective technique for the long term storage of SHINE reagents (*24*). Here, we utilize a lyophilization process in which monodisperse droplets are produced, flash-frozen in liquid nitrogen, lyophilized in strip tubes, and then placed in reaction chambers. Relative to previous SHINE lyophilization formulations, we find that by doubling the concentration of reagents and therefore halving the quantity of water for sublimation, the effectiveness of the lyophilization process is improved and the subsequent lyophilized pellets are more compact and much easier to transfer to reaction chambers. It is likely that reducing the water volume in the droplet minimizes deformation of the pellet as ice is sublimated, and reduces the surface area of the pellet exposed to the atmosphere, limiting hygroscopic effects. The lyophilized formulation utilizes cryoprotectants (sucrose) and excludes certain potentially destabilizing reagents (KCl, MgOAc, PEG) from the master mix (*24*). For this device, we only use sucrose, as mannitol, which has been used in previous SHINE formulations, inhibits the dissolution of the pellet. We observed that the formulation with mannitol and sucrose took several minutes to dissolve, whereas the formulation with just sucrose dissolved within seconds. With these alterations, the SONATA droplet lyophilization formulation retains its sensitivity when compared to the SHINE method as tested with Mtb synthetic targets **(Fig. 2D)**.

Combining the SONATA droplet lyophilization method with the on-chip magnetic mixing process, we demonstrate the single-plex detection of 10^4^ cp/μl of Mtb synthetic target on-chip (**Fig. 2E**,**F**). This shows that SHINE can be performed in a microfluidic device using lyophilized reagents.

### Sensitivity and specificity

To proceed with multiplexed on-chip detection, we chose the valve geometry for several reasons. The valve geometry is spatially compact, it allows for a chamber sealing step, and the serial filling process enables an easier containment of excess sample fluid. We made a few alterations to the original design to enhance these features. We find that the addition of a design feature that we call the ‘antechamber,’ a small chamber at the base of the inlet channel, acts as an additional surface tension valve, improving the sealing of the valve geometry using both oil and air. We also incorporate an on-chip reservoir, which is used to contain the excess sample fluid injected into the chip and displaced during the sealing process **(Fig. 3A)**. This reduces the risk of the user coming into contact with sample fluid. Also, we chose the air sealing method as it was found to be more robust than the oil sealing method throughout the incubation process.

**Figure 3.**
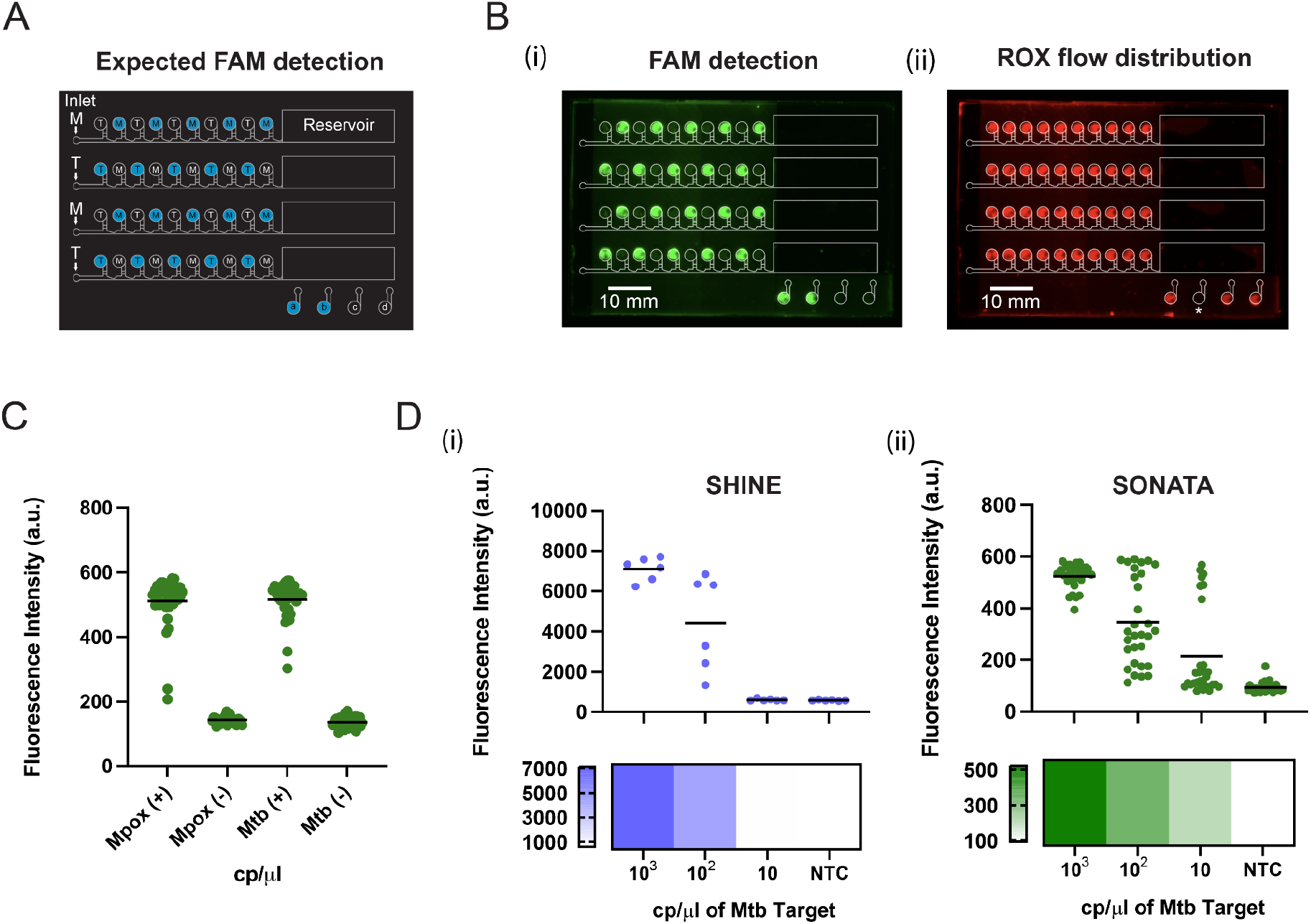
Multiplexed on-chip nucleic acid detection is sensitive and specific. **(A)** Schematic shows placement of Mpox (M) and Mtb (T) assays in each of the 4 independent channels on the chip such that all odd chambers contain Mtb and all even chambers contain Mpox guide sets. Chambers labeled in blue signify expected fluorescence based on the injection of the corresponding target in that channel. Control chambers in the bottom right corner correspond to (a) a 10^4^ cp/μl Mtb target control, (b) an always-on FAM reporter control, (c) a no MgOAc control, and (d) an NTC control. **(B) (i)** FAM fluorescent image of the chip at 60 minutes shows the expected staggered checkerboard pattern and the expected fluorescence in the two leftmost control chambers. **(B) (ii)** ROX fluorescent image of the chip at 60 minutes corresponding to the experiment in **(i)** shows even flow distribution. ROX reporter was not included in the always-on FAM control chamber labeled with an asterisk. The dark dot in each reaction chamber is the mixing bead. **(C)** Quantification of specificity data across four chips and two experiments showing the on-target (+) and off-target (-) fluorescence. The horizontal line represents the mean of n = 40 technical replicates. **(D)** Comparison of SHINE and SONATA sensitivity for different concentrations of Mtb synthetic target. The fluorescence for each method was measured by a different imaging system as described in the methods section leading to differences in the absolute fluorescence. **(i)** Performance at 60 min for standard SHINE. The horizontal line represents the mean of n = 6 technical replicates across two experiments. **(ii)** Performance at 60 min for SONATA. The horizontal line represents the mean of n = 30 technical replicates, corresponding to two experiments and 3 chips.

To validate that SONATA, incorporating on-chip mixing and lyophilization, performs comparably to standard SHINE in terms of sensitivity and specificity, we design a chip with four valve channels leading each to their own reservoir **(Fig. 3A)**. To demonstrate specificity, we distribute two different master mix formulations in lyophilized pellet form in a checkerboard pattern such that a pellet targeting Mtb is placed in all the even chambers and a pellet targeting Mpox is placed in all the odd chambers. Mpox and Mtb are distinct targets that should have no cross-talk at the sequence level. We use a high concentration of 10^4^ cp/μl of target input, as this stringently tests the performance of the chip in terms of specificity, as a high concentration of target makes evident any potential cross-talk between chambers. We find that during the incubation period, condensation in the central channel can lead to connections between chambers that allow for cross-talk. In order to prevent this, the chip needs to be oriented vertically during incubation, with the reservoir oriented downwards to allow gravitational forces to clear the central channel of condensation. Generally in microfluidic systems surface tension forces dominate over gravitational forces due to the small length scales (*34*). However, as the channel dimensions are on the larger side for our device (800 x 300 μm at their largest), gravitational forces can play some role.

After instituting these design features and injecting Mpox synthetic target in the first and third channel and Mtb synthetic target in the second and fourth channel and sealing the chip with a secondary air sealing step, we see the expected staggered checkerboard pattern of fluorescence as shown in **Fig. 3Bi**. This demonstrates that there is no cross-talk between reaction chambers. We also place ROX dye in the sample fluid to show that the flow distribution into each reaction chamber is uniform **(Fig. 3Bii)**. The performance of the device in terms of specificity is validated across two experiments and four chips in **Fig. 3C**.

To test the sensitivity of the device, we make a direct comparison to off-chip SHINE using the assay targeting Mtb. We make the two mastermixes in parallel according to the SHINE and SONATA lyophilization procedures. We image according to each procedure as described in the methods section, which leads to a distinction in absolute fluorescence levels. We see a similar performance in sensitivity for both methods **(Fig. 3D)**, which suggests that we can effectively multiplex SHINE on-chip without compromising sensitivity.

### Towards greater multiplexing

We sought to increase the level of multiplexing by an order of magnitude to enable deployable and highly multiplexed nucleic acid detection. To achieve this goal, it is necessary to make some key modifications to the valve geometry and to the SONATA procedure. To have such a large, integrated system entirely in series requires a relatively high input pressure, which could rupture the hydrophobic membrane used to seal the chip. To reduce the necessary applied pressure for flow distribution, we develop a 7 by 16 system, i.e., 112 chambers. The principle of this geometry is explained in the methods section. This hybrid design effectively combines the valve and comb geometries as the resistance-adjusted comb geometry is used to distribute the fluid to a set of valve geometries **(Fig. 4A)**.

**Figure 4.**
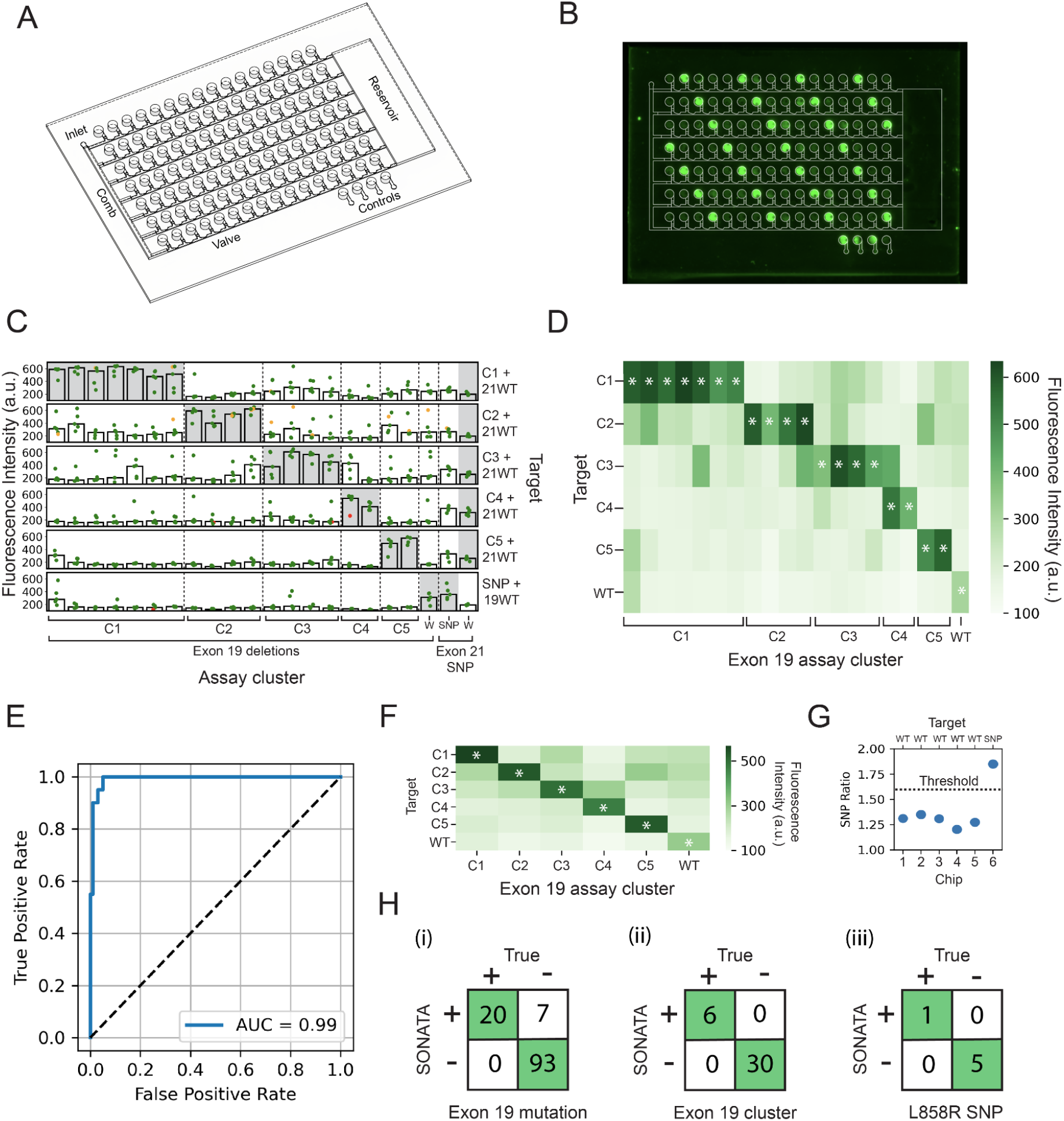
A hybrid 112-chamber configuration enables mutation discrimination. **(A)** The schematic shows a 3D representation of the hybrid 112-chamber geometry as a combination of the comb and valve geometries, a reservoir to contain excess fluid, and the inclusion of an NTC, no MgOAc, always-on and a 10^4^ cp/μl target control. **(B)** FAM fluorescent image of the 112-chamber hybrid geometry at 60 min shows the expected tiger stripes pattern. Mtb guide sets were placed along five diagonal stripes and Mpox guide sets were placed in all other chambers. Mtb synthetic target at 10^4^ cp/μl was injected into the chip. The small dark dot in each reaction chamber is the mixing bead. Control chambers in the bottom right corner correspond to a 10^4^ cp/μl Mtb target control, an always-on FAM reporter control, a no MgOAc control, and an NTC control. The no MgOAc control chamber was accidentally filled with the 10^4^ cp/μl Mtb target control due to pipetting error. **(C)** FAM fluorescence quantification at 60 minutes of 6 chips where each row represents a distinct chip and set of synthetic targets injected into that chip. The chip’s reaction chambers (110 out of 112 chambers) were pre-loaded with 22 different assays with 5 chambers per assay, targeting EGFR mutations associated with NSCLC. Specific assays and targets correspond to the order of sequences in **Table S2.** Chips 1 through 5 were injected with the corresponding cluster of synthetic targets, C1 through C5 of exon 19 deletion mutant sequences as well as the exon 21 WT sequence. Exon 19 deletions were clustered into pools based on sequence specificity as described in the methods section. Chip 6 was injected with the exon 19 WT sequence as well as the exon 21 L858R mutant sequence. All targets were at a concentration of 5×10^3^ cp/μl individually before pooling. Green dots signify the fluorescence of individual chambers, orange dots signify the fluorescence of chambers which were manually annotated to have high FAM fluorescence in the channel connecting to a neighboring chamber (suggesting potential cross-contamination), red dots signify chambers that were automatically determined to have low ROX fluorescence. The details of this processing are described in the methods section. These problematic chambers were not excluded from later processing. The bars represent the median of n = 5 technical replicates.The gray shading indicates the expected fluorescent results (SNP detection is determined as a ratio of SNP to WT fluorescence). **(D)** Heatmap of the median fluorescence at 60 minutes shows data from **(C)** with only exon 19 clusters and WT. Asterisks show the expected fluorescent results. **(E)** Receiver operator curve (ROC) shows the true positive and false positive rate across all 6 chips at the assay level for the exon 19 deletions and exon 19 WT assays. For ROC analysis, raw fluorescence was max-normalized within each chip, excluding the L858R assays, to account for the different overall target concentration in each chip due to pooled target injection. **(F)** Heatmap of the mean of the median fluorescence of each assay in each exon 19 cluster. **(G)** Dot-plot for the L858R SNP showing the ratio of the SNP to WT assay fluorescence for each chip with each target. The threshold of 1.57 (horizontal dotted line) was used to determine the presence of the mutation. This threshold was pre-determined based on a previous on-chip experiment as described in methods. **(H)** Concordance of SONATA and the known mutation presence at **(i)**, the individual assay level, **(ii)**, the cluster level and, **(iii)**, the L858R SNP across all 6 chips. **(i)** The threshold for detection was taken as 2-fold of the mean of all the NTC on-chip controls. In addition to the individual assay fluorescence, the NTC on-chip controls were first max-normalized within each chip before determining the mean fluorescence. **(ii)** The cluster was assigned based on the highest mean fluorescence of each cluster for each chip. **(iii)** The SNP presence was determined based on the raw fluorescence ratio of the SNP to WT crRNA and its value relative to the threshold shown in **G**.

To enable reliable air sealing across this 112-chamber system, the comb geometry needed modification. In an earlier prototype where the inlet was centrally located relative to the comb manifold and the direction of injection was parallel to the valve geometries, sealing was compromised as only the channels closest to the inlet were sealed fully **(Fig. S4)**. This design performed well, but for higher degrees of multiplexing we cannot tolerate a high failure rate. Thus, we redesigned the comb manifold so that the direction of injection in the main channel is perpendicular to that of the valve geometries, allowing for reliable sealing across the entire chip **(Fig. 4A)**.

To simplify the device construction, we develop an on-chip lyophilization method in contrast to the off-chip droplet lyophilization method that was used for the smaller manifolds. In this method, the chip dimensions are slightly enlarged to be compatible with a multichannel pipette, such that each tip aligns with alternating chambers. The SONATA master mix is pipetted into the reaction chambers and then the entire chip is flash frozen in liquid nitrogen and placed in a lyophilizer. We find that it is necessary to flash freeze the chip directly in liquid nitrogen to produce a sufficiently compact lyophilized cake. In addition, the ‘antechamber’ surface tension barrier is crucial to prevent master mix flowing out of the reaction chambers after the initial pipette distribution.

Incorporating these design elements, we achieve reliable flow distribution and sealing for the 112-chamber geometry as shown in **Fig. S3B**. The flow distribution and sealing works as follows. The comb geometry first distributes an even flow rate to each of the tributary valve geometries. Next, the first chamber in all the valve geometries fill in parallel **(Fig. S3D)**. Once the first set of chambers has filled, the valves burst and the next set of chambers fill. This process occurs in less than a minute of injection across the entire chip. In a subsequent step, injection of air into the chip via a squeeze bottle seals off all the chambers as independent reaction environments.

In order to validate the performance of the 112-chamber chip, we design an experiment in which Mtb master mix is distributed along a set of diagonals and all the remaining chambers are filled with Mpox master mix as shown in **Fig. S3A**, and the chip is lyophilized. After injecting the Mtb target at 10^4^ cp/μl into the chip and sealing, we see the expected tiger stripe pattern emerge after 60 minutes of incubation **(Fig. 4B)**. This tests and demonstrates specificity and assay performance across the entire 112-chamber geometry.

To further demonstrate the performance of SONATA, we seek to apply the technology to the detection of EGFR mutations implicated in NSCLC. Lung cancer is the leading global cause of cancer-related mortality and 85% of lung cancer cases are NSCLC (*35, 36*). The most common mutations in the EGFR gene are the exon 19 deletions and the exon 21 L858R point mutation (CTG to CGG), which account for approximately 90% of EGFR mutations in NSCLC (*37*). These mutations are of critical clinical importance, as they predict sensitivity to tyrosine kinase inhibitors (TKIs) and directly inform first-line treatment decisions (*37*). Despite their significance, a quarter of patients in Europe and North America begin treatment without knowing their mutational status, potentially undermining therapeutic effectiveness *(38, 39)*. This deficiency is even more pronounced in resource-limited settings: for example, a retrospective review from Ghana reported that only 17% of NSCLC cases underwent any biomarker testing at all (*40*). These gaps underscore a pressing need for accessible approaches to mutational characterization in NSCLC.

To develop our EGFR mutation panel, we chose to include all the exon 19 deletions in the Clinvar database corresponding to the two most common classes: the E746-A750 and L747-based deletions (*41*). The design process for crRNAs and primers are described in the methods section. To detect the L858R mutation, we designed crRNAs targeting both the SNP and the WT sequences. In addition, we employed crRNA-complementary DNA occluders, which enhance specificity by introducing a kinetic penalty to crRNA target binding (*42*). As shown in **Fig. S6B**, many of the exon 19 deletions are only a few mismatches apart in terms of crRNA-target pairs. We did not think that discriminating between these similar deletions was clinically relevant given that the phenotype, sensitivity to TKIs, and the resulting clinical outcomes are generally the same across exon 19 deletions (*43*). Thus, we chose to pool synthetic targets containing the exon 19 deletions into 5 pools for on-chip testing by sequence similarity as described in the methods assay design section. In addition, we predicted that pooling samples would be a more rigorous test of specificity by increasing the overall target concentration. We test these 5 pools of exon 19 deletions as well as the L858R SNP with 5 replicates each of 22 assays across six 112-chamber chips.

We evaluate the performance of SONATA at the individual assay level, the cluster level, and in terms of detection of the L858R SNP. At the assay level, as shown in **Fig. 4C** and**4D**, we see fluorescence of the expected assays with some off-target fluorescence, particularly for the chips with a greater number of pooled samples. As expected, higher overall target concentration leads to more off-target activity. For cluster 2, some of the off-target fluorescence can be ascribed to cross-contamination between chambers, which is noted in **Fig. 4C** by the orange dots. For cluster 3, the cross-talk with cluster 4 was predicted by the sequence analysis in **Fig. S6B**.

The results show that SONATA demonstrates highly robust detection. The receiver operator analysis yields an AUC of 0.99 **(Fig. 4E)**, a sensitivity of 100%, and a specificity of 93% based on a threshold of 2-fold the mean of the NTC controls **(Fig. 4Hi)**. At the cluster level, we report a sensitivity and specificity of 100% **(Fig. 4Hii)**. We detect the presence of the L858R SNP by taking a ratio of the SNP and WT fluorescence. As the SNP crRNA-occluder pair is more active than the WT crRNA-occluder pair this ratio is always greater than 1, but in all chips we still discriminate the presence of the SNP from the WT as shown in **Fig. 4G** based on a predetermined threshold of 1.57. This threshold was taken as the mean ratio of the SNP and WT condition in a previous on-chip experiment **(Fig. S3E)**.

## Discussion

Here we develop SONATA, a portable architecture capable of sensitive, specific, and highly-multiplexed nucleic acid detection, and demonstrate its performance for detecting pathogens and NSCLC mutations associated with drug susceptibility. SONATA has the potential to enable the on-site surveillance in animals and humans of clinically-relevant mutations and pathogens with ease, speed, and breadth, allowing for distributed multiplexed testing in resource-limited settings. This advance may ultimately facilitate the gathering of comprehensive and rich data at the point-of-care that was previously unattainable, informing patients, doctors, researchers, and policymakers.

Incorporating simple incubation methods, such as body heat or chemical heat packs, together with a phone-camera-based fluorescent readout, could further streamline SONATA’s equipment requirements (*25*). Additionally, integrating upstream or on-chip sample processing that eliminates the need for laboratory liquid handling would pave the way for true at-home operation. Ultimately, this work moves us closer to a testing paradigm in which untrained users can perform highly multiplexed nucleic acid self-testing at-home.

## Methods

### General Reagents

Chemical reagents were acquired through Sigma Aldrich unless otherwise specified. Oligonucleotide sequences were ordered from Integrated DNA Technologies (IDT) and are listed in Supplementary tables 2 through 4.

### Chip Design

#### Electrical analogies for microfluidics

Taking the Hagen-Poiseuille equation for pressure-driven flow in a pipe, where *L* and are, respectively, the length and radius of the pipe, and *Q* is the flow rate, we can define a hydrodynamic resistance (*R*_*h*_) (*34, 44*).

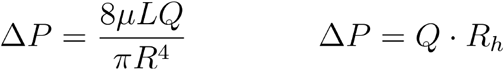

This introduces a useful analogy to Ohm’s law (Δ*V*=*I·R*_e_), where the electrical potential (Δ*V*) equates to the pressure gradient (Δ *P*), the current (*I*) equates to the volumetric flow rate (*Q*), and the electrical resistance (*R*_*e*_) equates to the hydrodynamic resistance (*R*_*h*_). In order to calculate and match hydrodynamic resistances across microfluidic systems, we can rely upon familiar relationships. If we take individual resistances to be, *R*_*m*_ where *m=*1,…,*N*,electrical analogies give expressions for effective resistances in series and in parallel (*34, 44*).

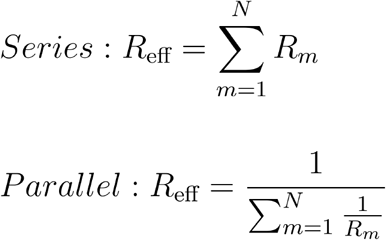

We can derive an expression for the hydrodynamic resistance in rectangular channels (as in our microfluidic system) starting from the Navier-Stokes equation and using cartesian coordinates where *w* is the width of the channel *h* and is the height (*34, 44*).

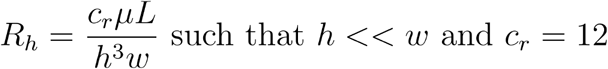

### Tree geometry

The tree geometry ensures an equal-path length from inlet to reaction chamber, which should give an equivalent hydrodynamic resistance. Given that the pressure drop across each path length is the same, we should expect an equal flow rate to each chamber (Δ*P=Q*.*R*_*h*_).

### Comb geometry

To fine tune the flow distribution among the secondary channels (**Fig. 1B**), we inserted at the base of each branch a short, elevated section whose length adjusts the hydraulic resistance of that branch. For instance, as illustrated in **Fig. 1B**, *Q* is the target flow rate entering each branch, and *R*_*n*_ denotes the hydraulic resistance of each branch. As the injected fluid travels along the primary channel toward this branch, the flow rate decreases stepwise because an amount *Q* is diverted into each preceding branch.

Moreover, the stream traverses a manifold segment with hydraulic resistance *R*^*^ . Accordingly, the total pressure drop to chamber 7 can be expressed as follows;

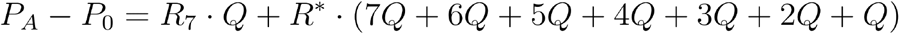

We can then generalize this expression for any chamber *n* where *N* represents the total number of chambers on one side of the manifold, which in this case would be 8, in which case,

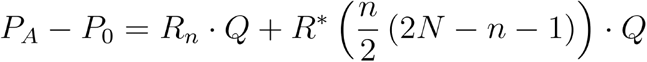

By substituting in the hydrodynamic resistance, equating this pressure drop to the total pressure drop for chamber 0, and setting the length of the elevated segment (*L*_0_) to 0, we derived a general expression for the length of each elevated segment (*L*_*n*_),

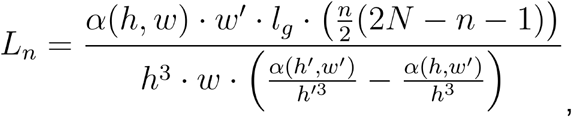

where *h* is the height of the primary channel and the elevated segment, *h*′ is the height of the secondary channel, *w* is the width of the primary channel, *w*′ is the width of the secondary channel and the elevated segment, and *l*_*g*_ is the length of each segment of the primary channel as shown in **Fig. 1B**. The expression *α*(*h, w*) is necessary to include in the hydrodynamic resistance when the height and width are similar,

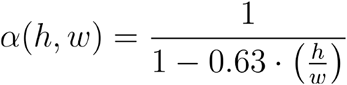

This general expression was used to determine the height of the elevated sections for both the comb and hybrid geometries.

### Valve geometry

In order to describe the effects of surface tension at these changes in geometry, particularly widening in either the height or width dimensions of a channel, we introduce the Young-Laplace equation, which describes the pressure difference across a curved interface due to surface tension,

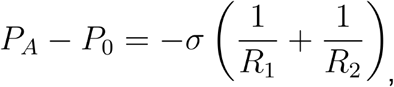

where *R*_1_ and *R*_2_ are the principal radii of curvature of the interface and *σ* is the surface tension (*45*).

Take a square channel with height *h* and width *w*. The fluid has a contact angle with the sidewalls (θ*s*), and with the top and bottom of the channel (θ*v*). We can recast the Young-Laplace equation in terms of these dynamic contact angles as shown in **Fig. S5B** (*28*), to find,

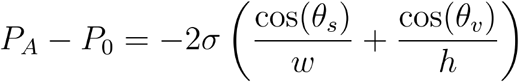

When the fluid approaches a widening channel, the interface is pinned, because the dynamic contact angle with the walls of the new channel,*θ*_*n*_, is less than the advancing contact angle (θ_*A*_) **(Fig. S5C)**. As the applied pressure increases, the interface bulges until *θ*_*n*_ reaches the advancing limit, *θ*_*A*_, at which point the meniscus depins (“bursts”). We define *β* as . *θ*_*A*_ − θ_*n*_ In terms of the contact angle with the original wall, bursting occurs when the contact angle is equivalent to *θ*_*A*_ +*β* = *θ*_*I*_. Given that 180° is the maximum contact angle that a liquid meniscus can attain, we modify our definition of θ_*I*_ to min(*θ*_*A*_*+ β*, 180°)(*28*), so that

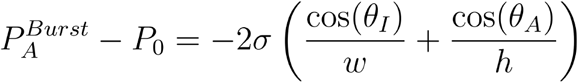

This gives us an expression for the burst pressure 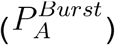 of a valve. As the burst pressure is inversely proportional to channel dimensions, by making valve 1 wider than valve 2, we can ensure that valve 1 bursts before valve 2 **(Fig. 1B)**.

### *Hybrid geometry* (112-chamber)

This hybrid geometry combines the comb and valve geometries as the resistance-adjusted comb geometry is used to distribute the fluid to a set of valve geometries. This combination of geometries is necessary to lower the pressure drop across the system. Taking our electrical analogies, we find that for a serial system the *Δ P* is equal to *N*·*QR*_*m*_, where is *N* the number of individual resistances (*R*_*m*_). On the other hand, for a parallel geometry, *Δ P* is equal to *QR*_*m*_ because although we increase the flow rate by a factor of *N*, this is counteracted by a reduction in the effective resistance by a factor of *N* . Hence, by using a 7× parallelized system, we are able to reduce the required *Δ P* by 7 relative to a purely serial geometry.

### Assay design

The Mpox guide sets were taken from the pre-designed assays on adapt.run for the Mpox virus. These designs were previously generated using ADAPT, a machine learning model trained to identify sensitivity guide target pairs, based on a multiple sequence alignment of available genomes and with the default parameters ((ID_M (4), ID_FRAC (0.01), BEST_N_TARGETS (10), PRIMER_LENGTH (30), PRIMER_COVER_FRAC (1), PRIMER_MISMATCHES (0) and MAX_PRIMERS_AT_SITE (unlimited)) (*46*). The top five (excluding the second due to secondary structure concerns) guides and primer sets with the highest predicted performance were chosen for testing. The reverse primer of the third guide set was adjusted from the original due to a high minimum △G of internal self-dimer formation. All the assays were found to show satisfactory performance as shown in **Fig. S6C**.

Guide set 3’ was used in this paper. For the Mtb assay, the IS6110C (also referred to as sg5) guide and primers and the IS6110A synthetic target were used (*47*) .

The exon 19 deletions and exon 21 L858R mutation make up 90% of EGFR mutations related to NSCLC (*37*). For the EGFR panel, we chose to include all the exon 19 deletions in the Clinvar database corresponding to the two most common classes (the E746-A750 and L747-based deletions) as well as the L858R point mutation (CTG to CGG) (*37, 41*). The exon 19 deletion crRNAs were designed such that the crRNA was centered at the start of the deletion to maximize the number of mismatches with the WT. If the mutation was an indel, the crRNAs were designed to be centered around the inserted sequence. For the exon 21 L858R SNP, three partially overlapping crRNAs were ordered so as to avoid a guanine directly 3’ to the target. Complementary (starting from the 5’ end of the guide) 23 nucleotide DNA oligos (occluders) with a Locked Nucleic Acid (LNA) at the eighth position were ordered to assist with mismatch detection. crRNA 2 was shown to have the best mismatch discrimination at 60 minutes as shown in **Fig. S6D**. A complementary WT crRNA was ordered with a corresponding 23 nucleotide occluder. It was later found that the sensitivity of the SNP crRNA was insufficient and so a 20 nucleotide occluder (shortened from the 5’ end of the occluder) was ordered to improve activity of this crRNA **(Fig. S6E)**. Exon 19 and exon 21 primers were designed following the TwistAMP guidelines for RPA primers (*48*). Primers were also screened for secondary structure using IDT Oligoanalyzer. As we aim to target genomic DNA, some primers extend into the introns. Synthetic targets for each mutation include the entire exon (exon 19 or 21) with 50 nucleotides of the introns appended to each end of the exon.

A T7 promoter sequence (5’-GAAATTAATACGACTCACTATAGGG-3’) was appended to the 5’ end of the forward primer and targets to allow for T7 RNA polymerase transcription of the RPA amplicon. The direct repeat (GAUUUAGACUACCCCAAAAACGAAGGGGACUAAAAC) was appended to the 5’ end of the guide sequence to yield the crRNA sequence.

#### Exon 19 deletion clustering

a specificity analysis of exon 19 deletions and synthetic targets was performed using NCBI BLAST. The BLAST output was filtered for alignments of 4 mismatches and below. The deletions were then clustered based on crRNA mismatch profiles as follows. The euclidean distance between each crRNA mismatch vector was hierarchically clustered, treating combinations without an alignment as 10 mismatches. The resulting dendrogram was thresholded to yield 5 clusters as shown in **Fig. S6A**.

### Assay formulation and lyophilization

#### Standard SHINE procedure

Both standard and lyophilized SHINE were performed as described in Bell et al with a few modifications. The SHINE formulation is as follows and was mixed in Ambion nuclease-free water: SHINE buffer (20 mM HEPES pH 8.0 with 60 mM KCl and 3.5% PEG-8000), 0.3 mM of each ribonucleotide triphosphate (rNTP) (NEB, cat. no. N0466L), 1 U µl−1 murine RNase inhibitor (NEB, cat. no.M0314L), 45 nM *Lwa*Cas13a purified as described below, 1 U/μl T7 RNAP (Biosearch Technologies Inc. (Lucigen, cat. no. NC2089983)), 62.5 nM FAM 6U quenched reporter (IDT), 45 nM of the crRNA, and 60 nM of pre-mixed RPA forward and reverse primers for all assays (except the c.2240_2254del which used 30 nM). The mixture of SHINE buffer, rNTPs, RNase inhibitor and nuclease-free water was used to resuspend TwistAmp Basic RPA pellets (TwistDx Limited, TABAS03KIT), whereby one pellet was resuspended for every 107.5 μl of final reaction volume, after which the rest of the mixture was added. MgOAc was added later at a final concentration of 14mM prior to adding target to yield the final master mix. Plate reader experiments were performed in triplicate with 10.5 ul reaction volumes in 384-well clear-bottom microplates (Greiner, cat. no. 788096). The reaction was then incubated at 37°C for up to 3 hours in an Agilent BioTek Cytation 5 microplate reader, as above. One-pot Cas13a assays were performed by mixing Cas13a master mix with target in a 9:1 ratio. The L858R SNP and WT crRNAs were pre-annealed with an occluder as described in the occluder annealing section.

For the lyophilized SHINE procedure, the SHINE buffer was substituted with just 20 mM HEPES (pH 8.0). Sucrose at 5% (w/v) and mannitol at 150 mM were added as cryoprotectants. Assays were aliquoted, flash frozen, and lyophilized at −20°C for 24 hours using a Freezone Triad Benchtop Freeze Dryer from Labconco. Aliquots were then vacuum sealed alongside a desiccant. Assays were resuspended in 3.5% PEG-8000, 60 mM KCl, and 14 mM MgOAc in Ambion nuclease-free water, aliquoted, and mixed with target. For the lyophilization comparison in **Fig. 2D**, the SHINE fluorescence was imaged via an Azure 600 imaging system as described in the Chip Testing section to make a direct comparison with the SONATA lyophilized pellets.

#### SONATA lyophilization procedure

The master mix for lyophilization was initially produced at 2X the final on-chip concentration to reduce the deformation of the pellet during the lyophilization process. The concentration of the pre-lyophilization formulation is as follows: 40 mM HEPES (pH 8.0), Sucrose (10% w/v), 0.6 mM of each ribonucleotide triphosphate (rNTP) (NEB, cat. no. N0466L), 2 U µl−1 murine RNase inhibitor (NEB, cat. no. M0314L), TwistAMP basic kit RPA pellets (1 pellet per unit reaction of volume 53.75 μl), purified *Lwa*Cas13a at 90 nM, 2 U/μl T7 RNAP (Biosearch Technologies Inc. (Lucigen, cat. no. NC2089983)), 125 nM PolyU Quenched FAM Reporter, 90nM Cas13 crRNA, 120 nM of pre-mixed RPA forward and reverse primers for all assays (except the c.2240_2254del which used 60 nM), and Ambion nuclease-free water. PEG 8000, MgOAc and KCl were excluded from the master mix and later added as part of the sample and resuspension buffer mixture. The L858R SNP and WT crRNAs were pre-annealed with an occluder as described in the occluder annealing section.

#### Off-chip lyophilization

After production of the master mix, droplets were flash frozen with a syringe pump (Harvard Apparatus 11 Elite Infusion) set at a flow rate of 60 ul/min oriented vertically over a liquid nitrogen bath. A 27 gauge needle tip (McMaster-Carr, cat. no. 75165A688) and a 1 ml syringe (Becton Dickinson, cat. no. 309628) was used to generate droplets of approximately 5.25 μl in volume. This is equivalent to 10.5 μl of master mix at the on-chip reaction concentration, which corresponds to the 11 μl chamber size accounting for the 0.5 μl mixing bead volume. Flash frozen droplets were transferred to strip tubes with plastic tweezers (McMaster-Carr, cat. no. 7003A35). Strip tubes were lyophilized in a cooling chamber (Electron Microscopy Sciences, cat. no.61951-10). After lyophilization they were stored at room temperature and used for one experiment within the same day.

#### On-chip lyophilization

Master mix was pipetted directly into the chip using a multichannel pipette. To assist with this process the chip was oriented upside-down on a tilted lab stand. This is to ensure that the master mix is pipetted into the opposite side of the inlet of each chamber to prevent potential clogging. Pipetting was performed slowly and only to the first stop to prevent bubble formation, which can compromise the lyophilization process. The chip was then slowly submerged in a liquid nitrogen bath for 10 seconds.

For both methods, the strip tubes or chips were transferred to the lyophilizer on dry ice and lyophilized for 24 hours in a Labconco Freezone Triad freeze dryer with the shelf temperature set to -20°C.

#### Occluder Annealing

crRNAs were pre-annealed to occluders in a mixture of 70mM KCl, 1uM crRNA and 10uM occluder in an Eppendorf Mastercycler X50 thermal cycler put through an annealing cycle consisting of a melting step at 85 ºC for three minutes followed by gradual cooling to 10 ºC at 0.1 ºC/sec and final cooling to 4 ºC.

### Chip construction

Chip geometries were designed using Autodesk Fusion 360, and CNC milling paths were created with FreeCAD. Molds were milled in polymethyl methacrylate (PMMA) using the Bantam Tools Desktop CNC Milling Machine. Polydimethylsiloxane (PDMS; Sylgard 184) with a 1:10 weight ratio of curing agent to elastomer base was mixed and defoamed in an AR-100 conditioning mixer (Thinky USA) and then poured into the mold, and baked in a convection oven at 70°C. After curing, the PDMS was removed from the mold and inlet/outlet holes were punched out using a 1.5 mm diameter biopsy punch (KAI Medical, BP-15F). A glass slide of the appropriate size (either 25 x 75 mm (Fisherbrand, cat. no. 12-550-A3) for the 16-chamber chips, 50 x 75 mm for the 4 x 10 plex chips (Fisherbrand, cat. no. 12-550C), and 102 x 83 mm (Ted Pella, cat. no. 260231) for the 112-chamber chips) was cleaned with ethanol and dried with compressed air. The PDMS layer and the glass slide were treated with air plasma (Harrick Plasma; 1 min) and immediately bonded to make a chip. Post surface treatments depended on channel geometry. For the tree and comb geometries, the chip was filled with polyethylene glycol (PEG) 200 for 25 minutes at 125°C to make hydrophilic surfaces (*49*). For the valve and hybrid geometries, the chip was treated with a hydrophobic coating (Aquapel, cat. no. 47100) for 5 minutes at room temperature. For the hybrid geometry, Aquapel was pipetted into each channel through a centrally located chamber to coat only the valve-based channels, not the comb manifold.

Aquapel is often counterfeited and should be purchased through certified distributors. Channels were then flushed continuously with 99.5% isopropyl alcohol (IPA) for 30 seconds and then dried with compressed air.

For the off-chip lyophilization method, lyophilized pellets were placed in reaction chambers using a 27 gauge needle tip (McMaster-Carr, cat. no. 75165A688; hereafter “27-gauge tip”)) Stir bars were made by cutting a 28 gauge Kanthal A-1 FeCrAl alloy wire to a length of roughly 1 mm, and placed in reaction chambers. Stir bars were exchanged with mixing beads (1 mm diameter steel ball bearings (uxcell, cat. no. B098SZFMRD) for more uniformity and ease of handling for the 4x10, and hybrid 112-chamber chips. Chips were sealed with either an optical adhesive film (Applied Biosystems) for the comb and tree geometries or low-density PTFE thread seal tape (Anti-seize Technology, cat. no. 26150) for the valve and hybrid geometries. As the optical film was not air permeable, it was punched at each chamber using a 27 gauge tip. Low-density PTFE thread seal tape was used to ensure sufficient air permeability. The tape was applied vertically across the entire chip with slight overlap between layers, leaving part of the reservoir uncovered to provide a visual cue for excess sample fluid.

Horizontal orientation was avoided, as overlapping layers in this direction can allow leakage during incubation. The tape was pressed by hand onto the PDMS without adhesive, and edges were trimmed. For the on-chip lyophilization method, any chambers in which the lyophilized cake appeared to clog the inlet channel potentially blocking flow into the chamber, were cleared with a 27 gauge tip dipped in ethanol and wiped with a Kimtech Kimwipe between clearing each new chamber.

### Chip testing

The resuspension buffer (60 mM KCl, 14mM MgOAc, 3.5% PEG 8000 and a 1:50 dilution of ROX Reference Dye (Thermo Fisher, cat. no. 12223012) at 25uM) was combined with the target in a 9:1 ratio to give the final resuspension solution.Resuspension solutions were injected into the chip using a 1 ml syringe (Becton Dickinson, cat. no. 309628) for the 16-chamber chip and 4 x10-chamber chip and a 2 ml syringe (Labfil, cat. no. C0001749) for the 112-chamber chip. Excess resuspension solution (200ul for the 4x10 chip and 1800ul for the 112-chamber chip) was injected with steady and low pressure into the chips and injection was stopped when fluid was visible entering the overflow reservoir. Though visualization is not necessary for injection, a lightbox (LitEnergy A4 LED Light Tracing Box) was used below chips during injection to visualize the filling of chambers through the PTFE membrane and confirm that the device was performing as expected. Chips were then sealed with a glass slide (of the same model as the base of the chip) covered with a layer of carbon double-sided tape (Nisshin-EM, cat. no. 7314), mixed via a hand driven rotation above a magnetic strip, and incubated at 37°C for times ranging from 60 to 90 minutes depending on the particular experiment. Mixing was performed for one minute by moving the chip in a circular motion above the magnetic strip and visually confirming that all the mixing beads are moving in response. The magnetic strip was composed of a series of neodymium bar magnets (FINDMAG, cat. no. B0B6PQPBP1). The 4x10-chamber and 112-chamber chips were placed vertically in a glass slide holder for incubation. Fluorescence was imaged through the glass bottom of the chip using the Azure 600 imaging system (Azure Biosystems) with a cy2 setting (excitation: 472 nm, emission: 513 nm) for FAM to analyze SHINE detection and a cy5 setting (excitation: 628 nm, emission: 684 nm) for ROX to analyze flow distribution. These settings were chosen to minimize cross-talk between the FAM and ROX channels. 500ms exposure was used for FAM fluorescent images and 2s exposure was used for ROX fluorescent images. Flow distribution was analyzed as above using just the resuspension buffer in **Fig. 1B,C**.

### Data analysis and figure generation

Fluorescent chip images shown are unprocessed. Analysis of chip fluorescence was performed using ImageJ (Version 1.53) where quantification of fluorescence intensity was taken as the integrated density without any background subtraction.The same set of ImageJ regions of interest (ROIs) were used to quantify chip fluorescence for each chip type to standardize analysis. The orange and red data points in **Fig. 4C** were annotated according to the following procedure. The red dots, which signify lower than expected ROX fluorescence in a chamber, were automatically annotated by an analysis script if the ROX fluorescence for a given chamber was less than 80% that of the mean ROX fluorescence of the No MgOAc, NTC and the 10^4^ cp/μl control chamber for that chip. The orange dots, which signify potential cross-contamination, were manually annotated if visible FAM fluorescence was seen in the channel connecting a chamber to another chamber. These chambers were still included in all downstream analysis. For receiver operator analysis in **Fig. 4**, the median fluorescences of each assay were normalized to the maximum median fluorescence for each chip (excluding the L858R SNP mutations which were processed separately). This was done to account for the different overall target concentrations in each chip. For the mutation level analysis of specificity and sensitivity, max-normalized fluorescences were also used. The NTC control of each chip was normalized to the maximum fluorescence within each chip and the threshold for detection was taken as 2-fold the mean of these normalized NTC values. For the cluster level analysis of sensitivity and specificity, the cluster was assigned based on the highest mean fluorescence for each cluster, taking the mean of the median fluorescence of each assay within a cluster. For the SNP level analysis, the ratio of the raw fluorescence of the L858R SNP assay to the WT assay was taken for each chip. Detection was determined based on whether the ratio exceeded the predetermined threshold of 1.57. This threshold was taken as the mean ratio of the SNP and WT condition in a previous on-chip experiment used for calibration.

#### Mixing analysis

9 μl of a 100% glycerol solution followed by 1 μl of ROX dye was pipetted into the reaction chamber as a stringent test of mixing in the highly viscous glycerol. The chamber was analyzed in a Leica SP5 confocal microscope and the resulting image sequence was processed in ImageJ to give a histogram of pixel intensity and the radial/axial slice images.

Schematics were made with Adobe Illustrator. Plots were generated with python seaborn (Version 0.12.2) and matplotlib (Version 3.6.2) packages as well as Prism (GraphPad, Version 10.4.1).

### *Lwa*Cas13a purification

The plasmid Plasmid pC013 - Twinstrep-SUMO-huLwCas13a (Addgene #90097) was transformed into Rosetta™ 2(DE3) Singles™ Competent Cells according to the manufacturer protocol (*16*). Single colonies were picked and inoculated in 30 mL of LB broth at 37C overnight. The culture was then scaled up by transferring the LB broth into 2L Terrific Broth and shaking at 37C and 200-250 rpm until the OD_600_ reached 0.6–0.8. The culture was then chilled on ice for 15–30 minutes before induction with 0.5 mM IPTG. It was then transferred to the shaker and incubated overnight at 18C.

The cells were pelleted by centrifugation at 4000 x g for 10 minutes and then resuspended in lysis buffer (20 mM Tris-HCl, 500 mM NaCl, 20 mM imidazole, 1 mM DTT, pH 8.0) supplemented with protease inhibitors (Thermo Scientific™ Pierce™ Protease Inhibitor Mini Tablets, EDTA-free, cat. no. A32955) and lysozyme (500μg/1ml) (Worthington Biochemical, cat. no. LS002880). Cells were lysed by sonication (Q500 Sonicator (Qsonica (Q500-110)) (10 s on, 10 s off, 50% power for 20 minutes) and the lysate was clarified by centrifugation at >20,000 × g for 1 hour at 4 °C. The supernatant was then filtered in a 45um syringe filter Cytiva, cat. no. 4654) before affinity purification in a Ni-NTA column via FPLC using wash **buffer A** (20 mM Tris-HCl, 500 mM NaCl, 1 mM DTT, pH 8.0) and elution **buffer B** (20 mM Tris-HCl, 500 mM NaCl, 300 mM imidazole, 1 mM DTT, pH 8.0). Eluted fractions were pooled and dialyzed overnight in SUMO cleavage buffer (30 mM Tris-HCl, 500 mM NaCl, 1 mM DTT, 0.15% NP-40, pH 8.0) in the presence of 300 µL of 1 mg/mL SUMO protease.

The sample was concentrated using a 50 kDa MWCO Amicon concentrator to half the volume, mixed 1:1 with cation exchange **buffer C** (20 mM HEPES, 250 mM NaCl, 1 mM DTT, 5% glycerol, pH 7.5), and purified on a 5 mL HiTrap SP HP column (cytiva (17115201)) via FPLC (Biorad NGC Quest 10 Plus Chromatography System, cat. no. 7880003). A salt gradient of 250 mM to 2M NaCl in elution **buffer D** (20 mM HEPES, 2000 mM NaCl, 1 mM DTT, 5% glycerol, pH 7.5) was used to elute bound proteins. SDS-PAGE (Genscript SurePAGE™, Bis-Tris, 10x8, 4-20%, 10 wells, cat. no. M00655) was used to check the purity of eluted fractions and those containing *Lwa*Cas13a were pooled accordingly.If sufficient purity was achieved, the protein was buffer-exchanged into storage **buffer E** (600 mM NaCl, 50 mM Tris-HCl, 5% glycerol, 2 mM DTT, pH 7.5), concentrated, aliquoted, and flash-frozen in liquid nitrogen for storage at ™80 °C. If purity was insufficient, the pooled protein was concentrated to 500 µL and subjected to size-exclusion chromatography (SEC) using a Superdex® 200 Increase 10/300 GL column (Cytiva, cat. no. 28990944). Fractions were appropriately pooled after SDS-PAGE (Genscript SurePAGE™, Bis-Tris, 10x8, 4-20%, 10 wells, cat. no. M00655) and the same buffer exchange and storage procedure was implemented.

## Acknowledgements

We thank Ben Larsen for assistance with occluder design; Alex Bell, Yong Ju, Owen Dunkley, Janine Nunes, and Ashwin Ramachandran for their valuable mentorship and insight; Long Nguyen for LwaCas13a purification; and Katie Wu for help with chip manufacturing. We are grateful to the Conway lab for access to the Freezone Triad Freeze Dryer, and to all members of the Myhrvold and Stone labs for their support.

## Author Contributions

S.K., R.S, H.A.S., and C.M. performed chip and experimental design. S.K.,Y.H., and S.H. performed assay design. S.K., R.S., Y.H., and I.M. performed experiments. S.K. and C.M analyzed data. All authors (S.K., R.S., Y.H., S.H., I.M., H.A.S., and C.M.) wrote the paper. C.M. and H.A.S. jointly supervised the work.

## Declaration of Interests

S.K., R.S., H.A.S., and C.M. are co-inventors on a patent application relating to this study.

## Funding

This work was supported by NIH R01 AI182281 (to C.M.), Centers for Disease Control and Prevention 75D30122C15113 (to C.M.), and the Princeton Catalysis Initiative (to C.M. and H.A.S.). R. Song was supported by the Princeton University Eric and Wendy Schmidt Transformative Technology Fund at Princeton University.

## Notes

### Summary of Updates

We have renamed mSHINE to SONATA to emphasize the generality of the microfluidic manifold. This resulted in a slight reorganization of the manuscript, including updates to the introduction and results sections.

