## Supplementary Information for "A handheld microfluidic manifold for massively multiplexed nucleic acid detection"

### Supplementary Figures

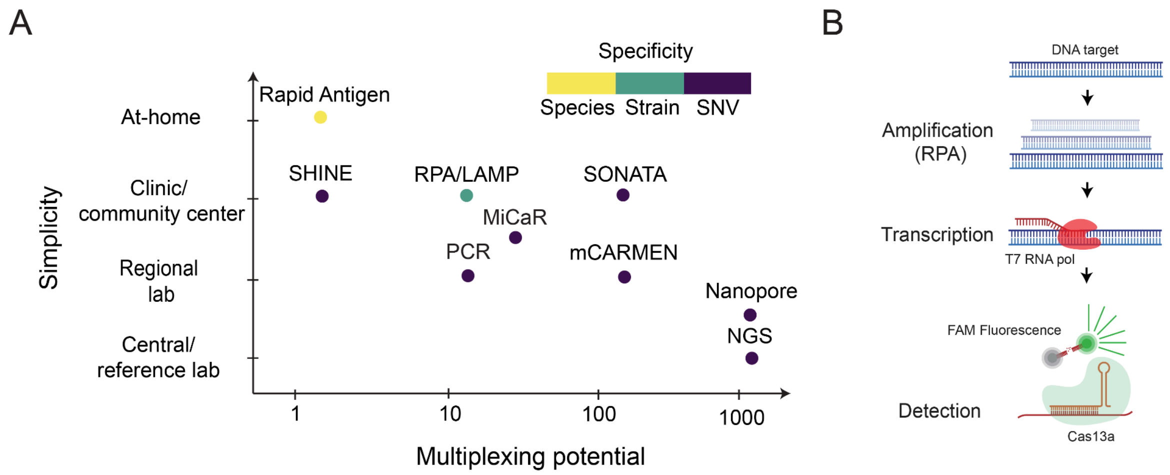


**Figure S1. (A)** Plot of multiplexing potential vs simplicity for nucleic acid detection and diagnostic technologies with the specificity annotated at the species, strain and single-nucleotide variation (SNV) level. Relevant citations are included in **Table S1. (B)** Biochemistry of the SHINE reaction, showing recombinase polymerase amplification (RPA), transcription by T7 RNA polymerase, and Cas13 cleavage of quenched FAM reporters upon the successful crRNA-mediated binding of the target sequence.

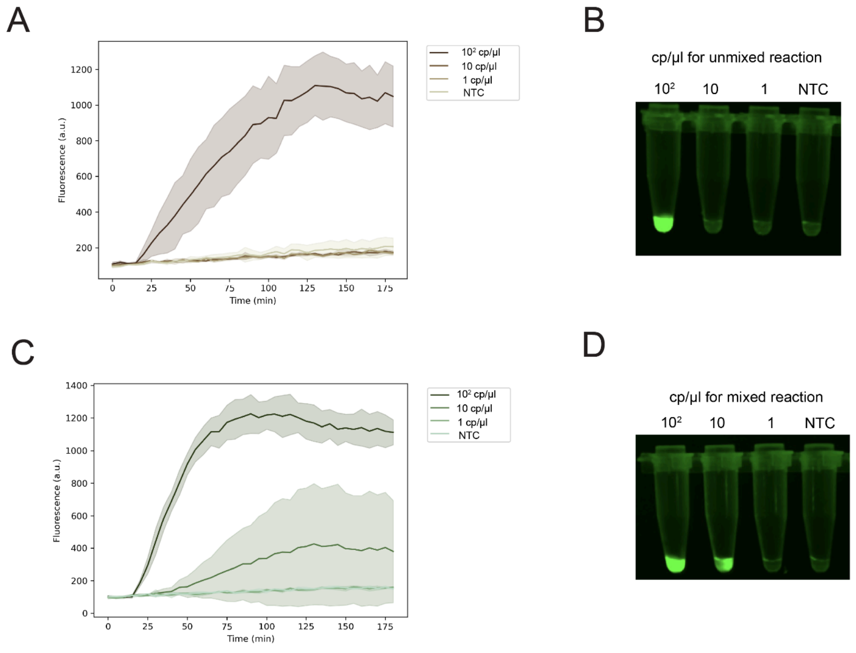


**Figure S2. Mixing importance for SHINE.** Performance of the SHINE reaction with non-lyophilized and lyophilized master mix for $10^2$, 10 and 1 cp/µl of mpox synthetic target. **(A)** Fluorescent kinetics of the SHINE reaction for a liquid master mix without vortex mixing prior to incubation. **(B)** FAM fluorescence at 60 minutes for the SHINE reaction with a lyophilized master mix without vortex mixing prior to incubation. **(C)** Fluorescent kinetics of the SHINE reaction for a liquid master mix with vortex mixing prior to incubation. **(D)** FAM fluorescence at 60 minutes for the SHINE reaction with a lyophilized master mix with vortex mixing prior to incubation.

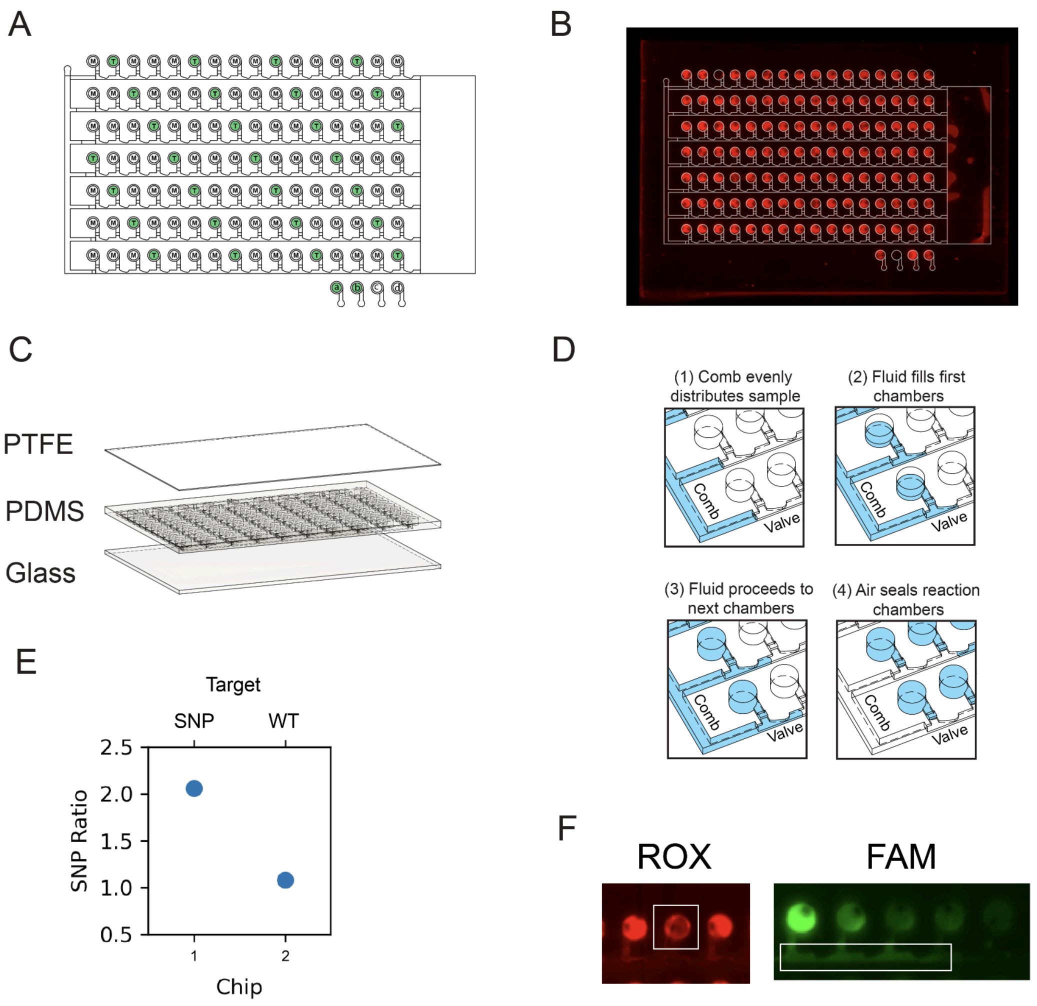


**Figure S3. Supporting data for Fig. 4. (A)** Schematic of the 112-chamber hybrid geometry shows the location of Mtb (T) and Mpox (M) guide sets. Mtb chambers are labelled in green as these are expected to be positive. Control chambers in the bottom right corner correspond to (a) a $10^4$ cp/µl Mtb target control, (b) an always-on FAM reporter control, (c) a no MgOAc control, and (d) an NTC control. **(B)** ROX fluorescent image of the experiment shown in **Fig. 4B**, demonstrating flow distribution. **(C)** Schematic shows the 3 layers of the hybrid geometry (PTFE, PDMS and glass). **(D)** Schematic shows how the combination of the comb and valve geometries enable flow distribution and sealing. (1-3) consists of the initial injection step, and (4) shows the result of the air sealing step. **(E)** Data generated from a 2-chip experiment where chip 1 was injected with the L858R SNP synthetic sequence and chip 2 was injected with the WT exon 21 sequence. Plot shows the ratio of the SNP crRNA to WT crRNA raw fluorescence. The mean of the two data points was used to generate the threshold of 1.57 for SNP detection used in **Fig. 4G**. **(F)** ROX and FAM fluorescent images of the chips corresponding to the experiment in **Fig. 4** show examples of a low-ROX chamber in the white box for the ROX image and a high-FAM channel in the white box for the FAM image.

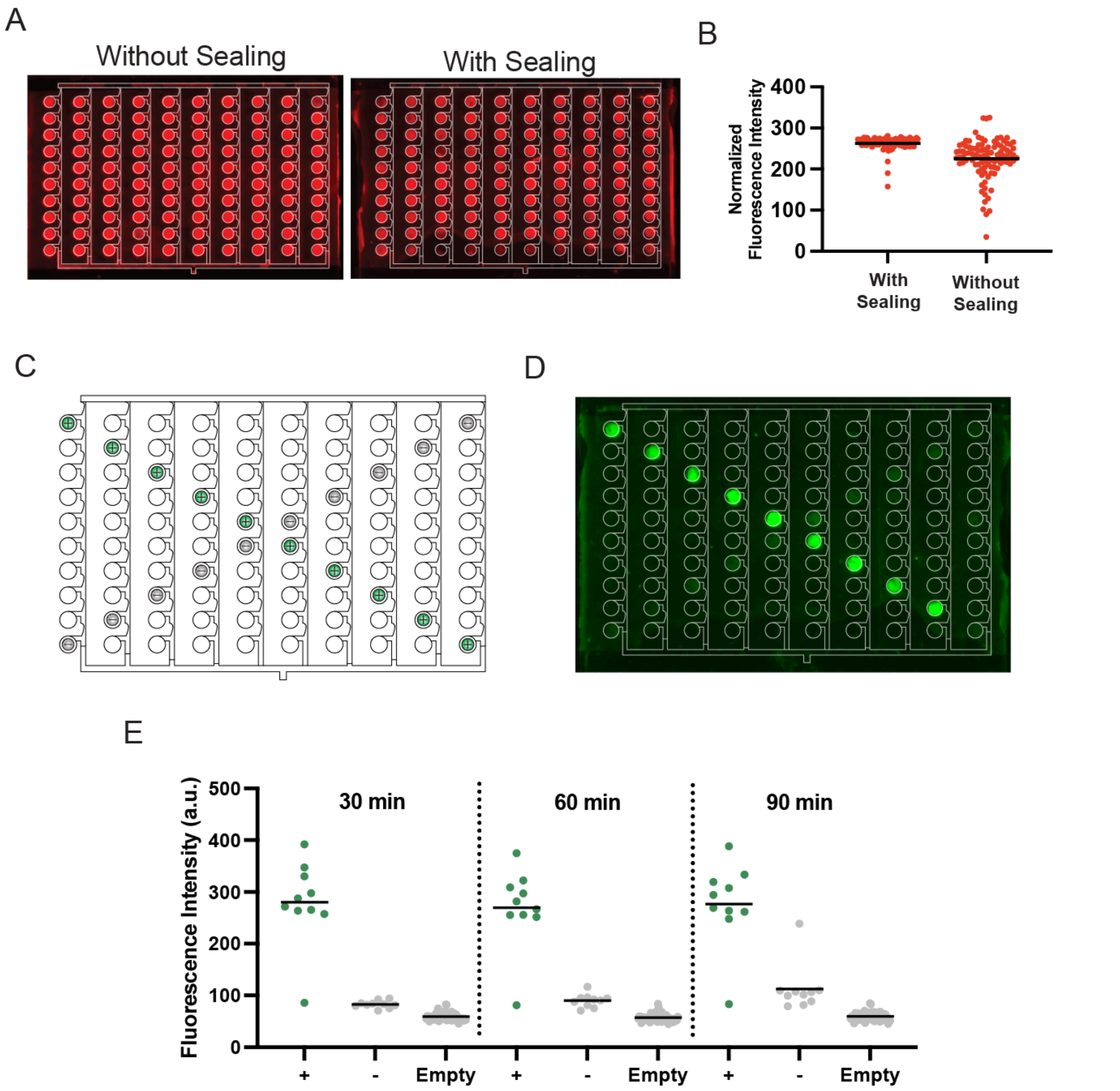


**Figure S4. Validation of the 100-chamber hybrid geometry. (A)** ROX fluorescent images show flow distribution for the hybrid geometry before and after the oil sealing step. For the image with sealing, the reduced fluorescence in the bottom row of chambers is caused by oil wetting the PTFE membrane and darkening it. Some chambers were partially compromised by oil. **(B)** Quantification of images in **A.** Background subtracted fluorescence was normalized by exposure time to account for a two-fold difference in exposure time. **(C)** Schematic of the hybrid geometry shows location of on-target (+) and off-target (-) guide sets with the remaining chambers not containing a lyophilized pellet. **(D)** FAM fluorescent image of chip shows the desired cross pattern after a 60 min incubation period with $10^5$ cp/µl of Mpox target. The bottom right chamber does not show any fluorescence, potentially due to the chamber being compromised by oil. **(E)** Quantification of FAM fluorescence in on-target (+), off-target (-) and empty chambers for 30, 60 and 90 minutes, corresponding to the experiment in **D.** The mean of n=10 technical replicates is shown for the on-target condition and off-target conditions. The empty condition shows the mean of n=80 technical replicates.

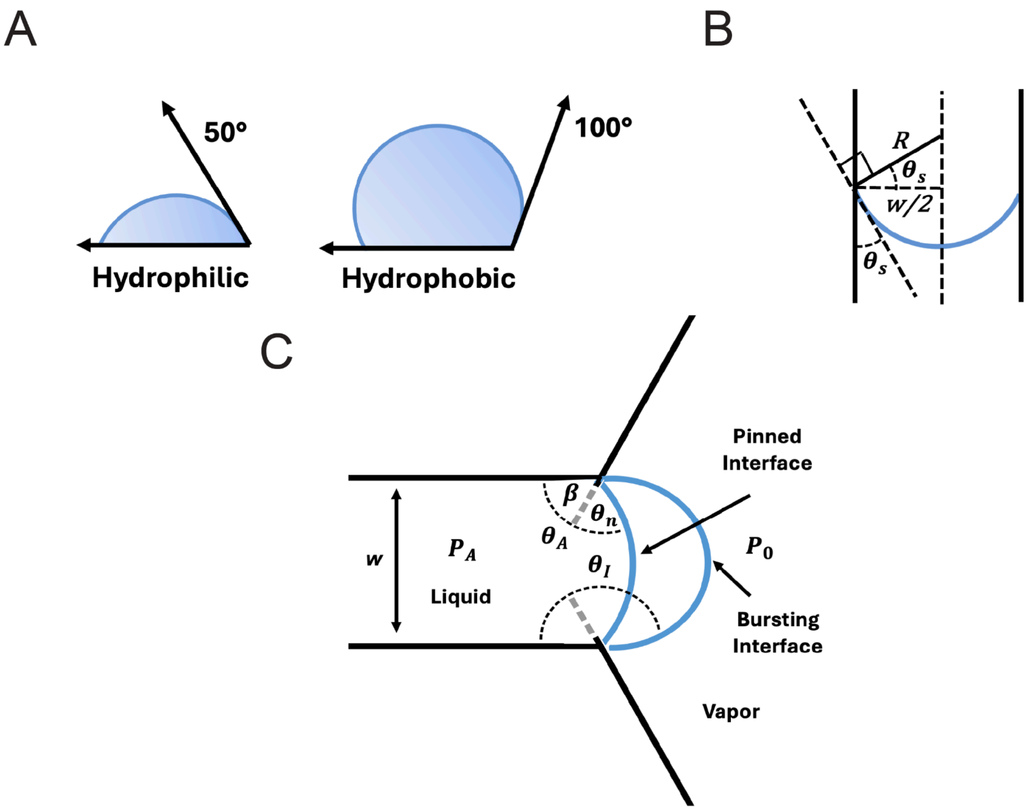


**Figure S5. Supporting schematics for the chip design section. (A)** Diagram of hydrophilic and hydrophobic equilibrium contact angles. **(B)** Diagram of a liquid with a hydrophilic contact angle flowing up a capillary, adapted from Pavuluri et al (*50*). The sketch shows how to express the Young-Laplace equation in terms of the contact angle with the sidewalls. From the diagram, R = w/2 cos($\theta_s$). **(C)** Diagram demonstrating the burst pressure, adapted from Cho et al (*28*).

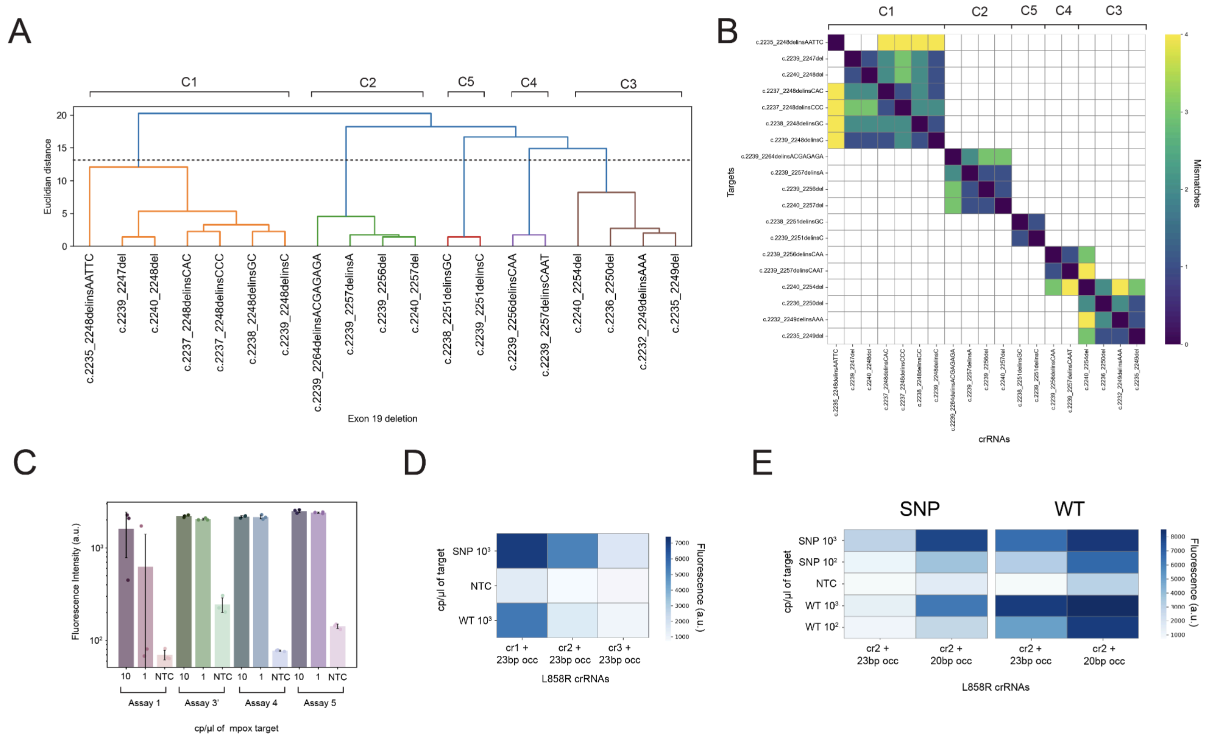


**Figure S6. Supporting data for the assay design section. (A)** Dendrogram of exon 19 deletions clustered by crRNA mismatch profiles as described in the methods assay design section. Cluster labels correspond to the clusters in **Fig. 4.** The order of specific deletions does not align with **Fig. 4**. The horizontal dotted line represents the threshold in euclidean distance used to define the five clusters. **(B)** Heatmap of NCBI BLAST local alignments between exon 19 deletion crRNAs and synthetic targets, color coded by the number of mismatches in these alignments. Corresponding clusters in **(A)** are labelled. **(C)** Barplot of the performance of the first, third, fourth and fifth ranking guide sets for Mpox generated by ADAPT as described in the methods assay design section. Plot shows the fluorescence for each assay for 10 and 1 cp/µl of Mpox synthetic target at 60 minutes. **(D)** Heatmap of the performance of the three initial crRNAs targeting the L858R SNP with 23bp occluders. crRNA2 (cr2) was chosen for further experiments due to robust discrimination from the WT. **(E)** Heatmap of the performance of crRNA2 (cr2) and its corresponding WT cr2 with 23bp and 20bp occluders for $10^3$ and $10^2$ cp/µl of SNP and WT targets.

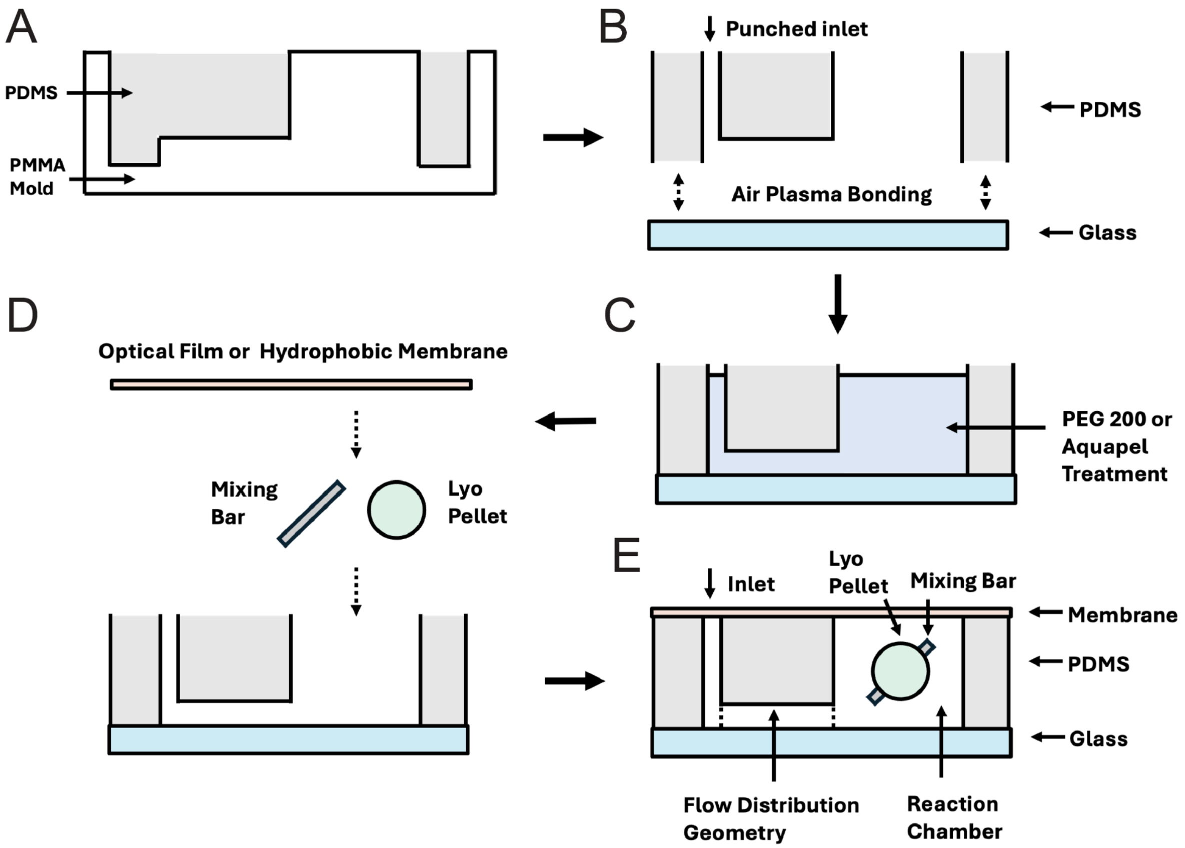


**Figure S7. Schematic showing the steps of chip construction**. **(A)** the curing of PDMS in the PMMA mold, **(B)** the outlet punching and air plasma bonding of PDMS to the glass slide, **(C)** the treatment of the chip with either PEG 200 (hydrophilic) or Aquapel (hydrophobic) coatings, **(D)** the addition of the lyophilized pellet, mixing bar and then sealing of the chip with either the optical film or hydrophobic membrane, and **(E)** the fully assembled chip.

**Table S1.** Comparison of nucleic acid detection and diagnostic technologies on sensitivity, specificity, accessibility, multiplexing and turnaround time.

| **Technology** | **Sensitivity** | **Specificity** | **Simplicity** | **Multiplexing** | **Turnaround Time** | **Reference** |
|---|---|---|---|---|---|---|
| PCR | very high | single-nucleotide | medium | 16 | 1-3 hours | (*51*, *52*) |
| NGS | very high | single-nucleotide | low | >1000 | 1-2 weeks | (*53*) |
| Nanopore Sequencing | very high | single-nucleotide | low-to-medium | >1000 | 7-9 hours | (*54*) |
| mCARMEN | very high | single-nucleotide | medium | >100 | 4 hours | (*26*) |
| MiCaR | high | single-nucleotide | medium-to-high | 30 | 40 min (on-chip assay) | (*27*) |
| Rapid Antigen Tests | medium | species | very high | 3 | 15 min | (*55*) |
| RPA/LAMP | high | strain | high | 16 | 30 min | (*56*) |
| SHINE | high | single-nucleotide | high | 2 | 1-2 hours | (*24*) |
| SONATA | high | single-nucleotide | high | >100 | 1-2 hours | this study |

**Table S2.** Synthetic target sequences of Mpox, Mtb and EGFR with e19 deletions organized by cluster.

| Name | Cluster | Target Sequence |
|---|---|---|
| Mpox Target | N/A | AAATAGATGTTAGTACAGTGTTATAAATGGATGAAGCATATTACTCTGGCA ACTTGGAATCAGTACTCGGGGATACGTGTCCGATATGCATACCGAACTC GCATCAATATCTCAATTAGTTATTGCCAAGATAGAAACTA |
| Mtb IS6110 | N/A | GAACCGGATCGATGTGTACTGAGATCCCCTATCCGTATGGTGGATAACG TCTTTCAGGTCGAGTACGCCTTCTTGTTGGCGGGTCCAGATGGCTTGC TCGATCGCGTCGAGGACCATGGAGGTGGCCATCGTGGAAGCGACCCG CCAGCCCAGGATCCTGCGAGCGTAGGCGTCGGTGACAAAGGCCACGT AGGCGAACCCTGCCCAGGTCGACACATAGGTGAGGTCTGCTACCCAC AGCCGGTTAGGTGCTGGTGGTCCGAAGCGGCGCTGGACGAGATCGG CGGGACGGGCTGTGGCCGGATCAGCGATCGTGGTCCTGCGGGCTTTG CCGCGGGTGGTCCCGGACAGGCCGAGTTTGGTCATCAGCCGTTCGAC GGTGCATCTGGCCACCTCGATGCCCTCACGGTTCAGGGTTAGCCACAC TTTGCGGGCACCGTAAACACCGTAGTTGGCGGCGTGGACGCGGCTGA TGTGCTCCTTGAGTTCGCCATCGCGCAGCTCGCGGCGGCTGGGCTCC CGGTTGATGTGGTCGTAGTAGGTCGATGGGGCGATCGGCACACCCAG CTCGGTCAGCTGTGTGCAGATCGACTCGACACCCCACCGCAAACCATC GGGGCCCTCGCGGTGGCCCTGATGATCGGCGATGAACCGGGTAATTA GCGTGCTGGCCGGTCGAGCTCGGCCGCGAAGAAAGCCGACGCGGTC TTTAAAATCGCGTTCGCCCTTCGCAATTCGGCGTTGTCCCGCCGCAAG CGCTTCAGCTCAGCGGATTCTTCGGTCGTGGTC |

| | | |
|---|---|---|
| c.2237_2248delinsCAC | C1 | GCACCATCTCACAATTGCCAGTTAACGTCTTCCTTCTCTCTCTGTCATAGGGACTCTGGATCCCAGAAGGTGAGAAAGTTAAAATTCCCGTCGCTATCAAGGCACCAACATCTCCGAAAGCCAACAAGGAAATCCTCGATGTGAGTTTCTGCTTTGCTGTGTGGGGGTCCATGGCTCTGAACCTCAGGCC |
| c.2237_2248delinsCCC | C1 | GCACCATCTCACAATTGCCAGTTAACGTCTTCCTTCTCTCTCTGTCATAGGGACTCTGGATCCCAGAAGGTGAGAAAGTTAAAATTCCCGTCGCTATCAAGGCCCCAACATCTCCGAAAGCCAACAAGGAAATCCTCGATGTGAGTTTCTGCTTTGCTGTGTGGGGGTCCATGGCTCTGAACCTCAGGCC |
| c.2239_2247del | C1 | GCACCATCTCACAATTGCCAGTTAACGTCTTCCTTCTCTCTCTGTCATAGGGACTCTGGATCCCAGAAGGTGAGAAAGTTAAAATTCCCGTCGCTATCAAGGAAGCAACATCTCCGAAAGCCAACAAGGAAATCCTCGATGTGAGTTTCTGCTTTGCGTGTGGGGGTCCATGGCTCTGAACCTCAGGCC |
| c.2240_2248del | C1 | GCACCATCTCACAATTGCCAGTTAACGTCTTCCTTCTCTCTCTGTCATAGGGACTCTGGATCCCAGAAGGTGAGAAAGTTAAAATTCCCGTCGCTATCAAGGAATCAACATCTCCGAAAGCCAACAAGGAAATCCTCGATGTGAGTTTCTGCTTTGCTGTGTGGGGGTCCATGGCTCTGAACCTCAGGCC |
| c.2238_2248delinsGC | C1 | GCACCATCTCACAATTGCCAGTTAACGTCTTCCTTCTCTCTCTGTCATAGGGACTCTGGATCCCAGAAGGTGAGAAAGTTAAAATTCCCGTCGCTATCAAGGAGCCAACATCTCCGAAAGCCAACAAGGAAATCCTCGATGTGAGTTTCTGCTTTGCTGTGTGGGGGTCCATGGCTCTGAACCTCAGGCC |
| c.2239_2248delinsC | C1 | GCACCATCTCACAATTGCCAGTTAACGTCTTCCTTCTCTCTCTGTCATAGGGACTCTGGATCCCAGAAGGTGAGAAAGTTAAAATTCCCGTCGCTATCAAGGAACCAACATCTCCGAAAGCCAACAAGGAAATCCTCGATGTGAGTTTCTGCTTTGCTGTGTGGGGGTCCATGGCTCTGAACCTCAGGCC |
| c.2235_2248delinsAATTC | C1 | GCACCATCTCACAATTGCCAGTTAACGTCTTCCTTCTCTCTCTGTCATAGGGACTCTGGATCCCAGAAGGTGAGAAAGTTAAAATTCCCGTCGCTATCAAAATTCCAACATCTCCGAAAGCCAACAAGGAAATCCTCGATGTGAGTTTCTGCTTTGCTGTGTGGGGGTCCATGGCTCTGAACCTCAGGCC |
| c.2239_2264delinsACGAGAGA | C2 | GCACCATCTCACAATTGCCAGTTAACGTCTTCCTTCTCTCTCTGTCATAGGGACTCTGGATCCCAGAAGGTGAGAAAGTTAAAATTCCCGTCGCTATCAAGGAAACGAGAGACAACAAGGAAATCCTCGATGTGAGTTTCTGCTTTGCTGTGTGGGGGTCCATGGCTCTGAACCTCAGGCC |
| c.2239_2257delinsA | C2 | GCACCATCTCACAATTGCCAGTTAACGTCTTCCTTCTCTCTCTGTCATAGGGACTCTGGATCCCAGAAGGTGAGAAAGTTAAAATTCCCGTCGCTATCAAGGAAACGAAAGCCAACAAGGAAATCCTCGATGTGAGTTTCTGCTTTGCTGTGTGGGGGTCCATGGCTCTGAACCTCAGGCC |
| c.2239_2256del | C2 | GCACCATCTCACAATTGCCAGTTAACGTCTTCCTTCTCTCTCTGTCATAGGGACTCTGGATCCCAGAAGGTGAGAAAGTTAAAATTCCCGTCGCTATCAAGGAACAACCGAAAGCCAACAAGGAAATCCTCGATGTGAGTTTCTGCTTTGCTGTGTGGGGGTCCATGGCTCTGAACCTCAGGCC |
| c.2240_2257del | C2 | GCACCATCTCACAATTGCCAGTTAACGTCTTCCTTCTCTCTCTGTCATAGGGACTCTGGATCCCAGAAGGTGAGAAAGTTAAAATTCCCGTCGCTATCAAGGAATCGAAAGCCAACAAGGAAATCCTCGATGTGAGTTTCTGCTTTGCTGTGTGGGGGTCCATGGCTCTGAACCTCAGGCC |
| c.2240_2254del | C3 | GCACCATCTCACAATTGCCAGTTAACGTCTTCCTTCTCTCTCTGTCATAGGGACTCTGGATCCCAGAAGGTGAGAAAGTTAAAATTCCCGTCGCTATCAAGGAATCTCCGAAAGCCAACAAGGAAATCCTCGATGTGAGTTTCTGCTTTGCTGTGTGGGGGTCCATGGCTCTGAACCTCAGGCC |

| | | |
|---|---|---|
| c.2232_2249delinsAAA | C3 | GCACCATCTCACAATTGCCAGTTAACGTCTTCCTTCTCTCTCTGTCATAG<br>GGACTCTGGATCCCAGAAGGTGAGAAAGTTAAAATTCCCGTCGCTATAA<br>AAACATCTCCGAAAGCCAACAAGGAAATCCTCGATGTGAGTTTCTGCTT<br>TGCTGTGTGGGGGTCCATGGCTCTGAACCTCA<br>GGCC |
| c.2235_2249del | C3 | GCACCATCTCACAATTGCCAGTTAACGTCTTCCTTCTCTCTCTGTCATAG<br>GGACTCTGGATCCCAGAAGGTGAGAAAGTTAAAATTCCCGTCGCTATCA<br>AAACATCTCCGAAAGCCAACAAGGAAATCCTCGATGTGAGTTTCTGCTT<br>TGCTGTGTGGGGGTCCATGGCTCTGAACCTCA<br>GGCC |
| c.2236_2250del | C3 | GCACCATCTCACAATTGCCAGTTAACGTCTTCCTTCTCTCTCTGTCATAG<br>GGACTCTGGATCCCAGAAGGTGAGAAAGTTAAAATTCCCGTCGCTATCA<br>AGACATCTCCGAAAGCCAACAAGGAAATCCTCGATGTGAGTTTCTGCTT<br>TGCTGTGTGGGGGTCCATGGCTCTGAACCTCAGGCC |
| c.2239_2256delinsCAA | C4 | GCACCATCTCACAATTGCCAGTTAACGTCTTCCTTCTCTCTCTGTCATAG<br>GGACTCTGGATCCCAGAAGGTGAGAAAGTTAAAATTCCCGTCGCTATCA<br>AGGAACAACCGAAAGCCAACAAGGAAATCCTCGATGTGAGTTTCTGCT<br>TTGCTGTGTGGGGGTCCATGGCTCTGAACCTCAGGCC |
| c.2239_2257delinsCAA<br>T | C4 | GCACCATCTCACAATTGCCAGTTAACGTCTTCCTTCTCTCTCTGTCATAG<br>GGACTCTGGATCCCAGAAGGTGAGAAAGTTAAAATTCCCGTCGCTATCA<br>AGGAACAATCGAAAGCCAACAAGGAAATCCTCGATGTGAGTTTCTGCTT<br>TGCTGTGTGGGGGTCCATGGCTCTGAACCTCAGGCC |
| c.2238_2251delinsGC | C5 | GCACCATCTCACAATTGCCAGTTAACGTCTTCCTTCTCTCTCTGTCATAG<br>GGACTCTGGATCCCAGAAGGTGAGAAAGTTAAAATTCCCGTCGCTATCA<br>AGGAGCCATCTCCGAAAGCCAACAAGGAAATCCTCGATGTGAGTTTCT<br>GCTTTGCTGTGTGGGGGTCCATGGCTCTGAACCTCAGGCC |
| c.2239_2251delinsC | C5 | GCACCATCTCACAATTGCCAGTTAACGTCTTCCTTCTCTCTCTGTCATAG<br>GGACTCTGGATCCCAGAAGGTGAGAAAGTTAAAATTCCCGTCGCTATCA<br>AGGAACCATCTCCGAAAGCCAACAAGGAAATCCTCGATGTGAGTTTCG<br>CTTTGCTGTGTGGGGGTCCATGGCTCTGAACCTCAGGCC |
| e19_WT | C6 | GCACCATCTCACAATTGCCAGTTAACGTCTTCCTTCTCTCTCTGTCATAG<br>GGACTCTGGATCCCAGAAGGTGAGAAAGTTAAAATTCCCGTCGCTATCA<br>AGGAATTAAGAGAAGCAACATCTCCGAAAGCCAACAAGGAAATCCTCG<br>ATGTGAGTTTCTGCTTTGCTGTGTGGGGGTCCATGGCTCTGAACCTCA<br>GGCC |
| >c.2573T>G (L858R<br>SNP) | N/A | CTTCTTCCCATGATGATCTGTCCCTCACAGCAGGGTCTTCTCTGTTTCA<br>GGGCATGAACTACTTGGAGGACCGTCGCTTGGTGCACCGCGACCTGG<br>CAGCCAGGAACGTACTGGTGAAAACACCGCAGCATGTCAAGATCACAG<br>ATTTTGGGCGGGCCAAACTGCTGGGTGCGGAAGAGAAAGAATACCATG<br>CAGAAGGAGGCAAAGTAAGGAGGTGGCTTTAGGTCAGCCAGCATTTTC<br>CTGACACCAGGGACCA |
| e21_WT | N/A | CTTCTTCCCATGATGATCTGTCCCTCACAGCAGGGTCTTCTCTGTTTCA<br>GGGCATGAACTACTTGGAGGACCGTCGCTTGGTGCACCGCGACCTGG<br>CAGCCAGGAACGTACTGGTGAAAACACCGCAGCATGTCAAGATCACAG<br>ATTTTGGGCTGGCCAAACTGCTGGGTGCGGAAGAGAAAGAATACCATG<br>CAGAAGGAGGCAAAGTAAGGAGGTGGCTTTAGGTCAGCCAGCATTTTC<br>CTGACACCAGGGACCA |

**Table S3.** crRNA sequences of Mpox, Mtb and EGFR with e19 deletions organized by cluster.

| Name | Cluster | crRNA Sequence |
|---|---|---|
| Mpox crRNA | N/A | GAUUUAGACUACCCCAAAAACGAAGGGGACUAAAACCUACAAGAGAGAGCUUGAUGAGACAACG |
| Mtb IS6110 C crRNA | N/A | GAUUUAGACUACCCCAAAAACGAAGGGGACUAAAACCUACGGUGUUUACGGUGCCCGCAAAGUG |
| c.2237_2248delinsCAC | C1 | GAUUUAGACUACCCCAAAAACGAAGGGGACUAAAACUCGGAGAUGUUGGUGCCUUGAUAGCGAC |
| c.2237_2248delinsCCC | C1 | GAUUUAGACUACCCCAAAAACGAAGGGGACUAAAACUCGGAGAUGUUGGGGCCUUGAUAGCGAC |
| c.2239_2247del | C1 | GAUUUAGACUACCCCAAAAACGAAGGGGACUAAAACUUUCGGAGAUGUUGCUUCCUUGAUAGCG |
| c.2240_2248del | C1 | GAUUUAGACUACCCCAAAAACGAAGGGGACUAAAACCUUUCGGAGAUGUUGAUUCCUUGAUAGC |
| c.2238_2248delinsGC | C1 | GAUUUAGACUACCCCAAAAACGAAGGGGACUAAAACUUCGGAGAUGUUGGCUCCUUGAUAGCGA |
| c.2239_2248delinsC | C1 | GAUUUAGACUACCCCAAAAACGAAGGGGACUAAAACUUCGGAGAUGUUGGUUCCUUGAUAGCGA |
| c.2235_2248delinsAATTC | C1 | GAUUUAGACUACCCCAAAAACGAAGGGGACUAAAACCGGAGAUGUUGGAAUUUUGAUAGCGACG |
| c.2239_2264delinsACGAGAGA | C2 | GAUUUAGACUACCCCAAAAACGAAGGGGACUAAAACUUCCUUGUUGUCUCUCGUUUCCUUGAUA |
| c.2239_2257delinsA | C2 | GAUUUAGACUACCCCAAAAACGAAGGGGACUAAAACUUGUUGGCUUUCGUUUCCUUGAUAGCGA |
| c.2239_2256del | C2 | GAUUUAGACUACCCCAAAAACGAAGGGGACUAAAACCUUGUUGGCUUUCGGUUCCUUGAUAGCG |
| c.2240_2257del | C2 | GAUUUAGACUACCCCAAAAACGAAGGGGACUAAAACCCUUGUUGGCUUUCGAUUCCUUGAUAGC |
| c.2240_2254del | C3 | GAUUUAGACUACCCCAAAAACGAAGGGGACUAAAACUGUUGGCUUUCGGAGAUUCCUUGAUAGC |
| c.2232_2249delinsAAA | C3 | GAUUUAGACUACCCCAAAAACGAAGGGGACUAAAACUUCGGAGAUGUUUUUAUAGCGACGGGAA |
| c.2235_2249del | C3 | GAUUUAGACUACCCCAAAAACGAAGGGGACUAAAACGCUUUCGGAGAUGUUUUGAUAGCGACGG |
| c.2236_2250del | C3 | GAUUUAGACUACCCCAAAAACGAAGGGGACUAAAACGGCUUUCGGAGAUGUCUUGAUAGCGACG |
| c.2239_2256delinsCAA | C4 | GAUUUAGACUACCCCAAAAACGAAGGGGACUAAAACGUUGGCUUUC |

| | | |
|---|---|---|
| | | GGUUGUUCCUUGAUAGCG |
| c.2239_2257delinsCAAT | C4 | GAUUUAGACUACCCCAAAAACGAAGGGGACUAAAACUGUUGGCUUUCGAUUGUUCCUUGAUAGC |
| c.2238_2251delinsGC | C5 | GAUUUAGACUACCCCAAAAACGAAGGGGACUAAAACGCUUUCGGAGAUGGCUCCUUGAUAGCGA |
| c.2239_2251delinsC | C5 | GAUUUAGACUACCCCAAAAACGAAGGGGACUAAAACGCUUUCGGAGAUGGUUCCUUGAUAGCGA |
| e19_WT | N/A | GAUUUAGACUACCCCAAAAACGAAGGGGACUAAAACCGGAGAUGUUGCUUCUCUUAAUUCCUUG |
| >c.2573T>G (L858R SNP)_cr1 | N/A | GAUUUAGACUACCCCAAAAACGAAGGGGACUAAAACUUUGGCCCGCCCAAAAUCUGUGAUCUUG |
| >c.2573T>G (L858R SNP)_cr2 | N/A | GAUUUAGACUACCCCAAAAACGAAGGGGACUAAAACCAGUUUGGCCCGCCCAAAAUCUGUGAUC |
| >c.2573T>G (L858R SNP)_cr3 | N/A | GAUUUAGACUACCCCAAAAACGAAGGGGACUAAAACCCCAGCAGUUUGGCCCGCCCAAAAUCUG |
| >c.2573T>G (L858R WT) | N/A | GAUUUAGACUACCCCAAAAACGAAGGGGACUAAAACCAGUUUGGCCAGCCCAAAAUCUGUGAUC |

**Table S4.** Additional miscellaneous sequences including primers, occluders and reporters.

| Name | Sequence |
|---|---|
| >c.2573T>G (L858R SNP) Occluder 23bp | CAGATTT+TGGGCGGGCCAAACTG |
| >c.2573T>G (L858R SNP) Occluder 20bp | ATTT+TGGGCGGGCCAAACTG |
| >c.2573T>G (L858R WT) Occluder 23bp | CAGATTT+TGGGCTGGCCAAACTG |
| >c.2573T>G (L858R WT) Occluder 20bp | ATTT+TGGGCTGGCCAAACTG |
| e19 del F Primer | GAAATTAATACGACTCACTATAGGGAACGTCTTCCTTCTCTCTCTGTCATAGGGACTCTG |
| e19 del R Primer | ATGGACCCCCACACAGCAAAGCAGAAACTCACAT |
| e21 L858R F Primer | GAAATTAATACGACTCACTATAGGGAGCCAGGAACGTACTGGTGAAAACACCGCAGCATG |
| e21 L858R R Primer | CTTTGCCTCCTTCTGCATGGTATTCTTTCTCTTCC |

| | |
|---|---|
| Mtb F Primer | GAAATTAATACGACTCACTATAGGGCCACCTCGATGCCCTCACGGTTCAGGGTTAGCCA |
| Mtb R Primer | GCGAACTCAAGGAGCACATCAGCCGCGTCCACGCCGCCAA |
| Mpox F Primer | GAAATTAATACGACTCACTATAGGGTAAATGGATGAAGCATATTACTCTGGCAAC |
| Mpox R Primer | TTGGCAATAACTAATTGAGATATTGATGCGAGTTCGGTAT |
| 6U-FAM-reporter | /56-FAM/rUrUrUrUrUrU/3IABkFQ/ |
